# Liver environment-imposed constraints diversify movement strategies of liver-localized CD8 T cells

**DOI:** 10.1101/2020.11.06.371690

**Authors:** Harshana Rajakaruna, James O’Connor, Ian A. Cockburn, Vitaly V. Ganusov

## Abstract

Pathogen-specific CD8 T cells face the problem of finding rare cells that present their cognate antigen either in the lymph node or infected tissue. While quantitative details of T cell movement strategies in some tissues such lymph nodes or skin have been relatively well characterized, we still lack quantitative understanding of T cell movement in many other important tissues such as the spleen, lung, liver, and gut. Furthermore, how environmental constraints influence movement patterns of T cells in tissues remains incompletely characterized. Liver is one of the major organs in mice and humans and is the target site for many pathogens such as hepatitis B and C viruses in humans and *Plasmodium* parasites in multiple mammalian species. We developed a protocol to generate stable numbers of liver-located CD8 T cells, used intravital microscopy to record movement patterns of CD8 T cells in livers of live mice, and analyzed these and previously published data using well-established statistical and computational methods. We show that in most of our experiments *Plasmodium*-specific liver-localized CD8 T cells perform correlated random walks characterized by transiently superdiffusive displacement with persistence times of 10-15 min that exceed those observed for T cells in lymph nodes. Liver-localized CD8 T cells typically crawl on the lumenal side of liver sinusoids (i.e., are in the blood); simulating T cell movement in digital structures derived from the liver sinusoids illustrates that liver structure alone is sufficient to explain the relatively long superdiffusive displacement of T cells. In experiments when CD8 T cells in the liver poorly attach to the sinusoids (e.g., one week post immunization with radiation-attenuated *Plasmodium* sporozoites) T cells also undergo Levy flights – large displacements occurring due to cell deattaching from the endothelium, floating with the blood flow, and re-attaching at another location. Our analysis, thus, provides quantitative details of movement patterns of liver-localized CD8 T cells and illustrates how structural and physiological details of the tissue may impact T cell movement patterns.

## Introduction

Search for resources is a fundamental property of many biological agents starting from individual cells to large animals. While different agents involve various search strategies, two types of search are of particular importance due to their fundamental nature: Brownian walks and Levy walks/flights [1]. Brownian walkers display short displacements, typically consistent with a thin-tailed distribution that has a finite mean and variance, with random turning angles; in Brownian walks mean squared displacement (MSD) increases linearly with time [1–4]. If a walker exhibits preference in movement direction similar to the previous movement direction, this results in correlated random walks during which MSD increases transiently faster than linearly with time and can be characterized by so-called persistence time [5–7]. In contrast, Levy walkers display both short and long displacements consistent with a fat-tailed distribution that has either finite mean and infinite variance or both infinite mean and variance; in Levy walks/flights MSD increases faster than linear with time [1, 3]. Various combinations of these two extremes such as composite correlated random walks, generalized Levy walks have been also studied (e.g., [8]). Understanding walk types of various biological agents has been a subject of many investigations and several techniques such as analysis of movement length distribution and MSD dynamics have emerged as relatively standard [7–17].

For adequate protection T cells of vertebrates survey tissues for infection by moving within tissues. Yet, specific movement strategies and impact of tissue environments on the movement patterns remain incomplete understood for most tissues in vertebrates such as mice. In lymph nodes (LNs) naive T cells seem to search randomly for antigen-presenting cells, acting as Brownian walkers undergoing short displacements in correlated random walks, although some role of attracting chemokines towards dendritic cells (DCs) expressing cognate antigen has been proposed [3, 6, 18–20]. In the skin, activated or memory CD8 T cells tend to perform Brownian walks [21], and in explanted murine lungs T cells display intermittent migration resulting in normal diffusion and a large variability in average velocities between individual cells although specific type of the walk (Brownian, Levy or a combination) has not been determined [22]. Interestingly, *Toxoplasma gondii*-specific activated CD8 T cells in murine brains undergo Levy walks although specific mechanisms (whether cell-intrinsic or environmentally driven) allowing such T cells to display rare long displacements have not been determined [8].

Liver is a large organ in mice and humans and is the target of multiple pathogens [23]. Apicomplexan parasite *Plasmodium*, the causative agents of malaria, infects the liver following inoculation of parasites by probing mosquitoes [24–26]. At this stage of the *Plasmodium* lifecycle the intracellular liver stage parasites are susceptible to clearance by vaccine-induced CD8 T cells. Remarkably, liver-localized CD8 T cells alone are capable of eliminating all liver stages and preventing clinical malaria [27, 28]. Given that a murine liver has about 10^8^ hepatocytes how exactly CD8 T cells locate all parasites within the 48 hour span of the parasite liver stages is unknown [29]. Furthermore, while we and others have documented movement of liver-localized CD8 T cells, the walk types employed by these cells and their potential underlying mechanisms remain incompletely understood [30–32].

Typical intravital imaging experiments allow to collect 3D coordinates of T cells in tissues over time, and with such data one can construct several movement characteristics to determine the type of walk cells exhibit [5–8, 12, 33]. Specifically, the rate of MSD increase with time, distributions of movement lengths and turning angles often allow to determine the walk type [7, 12, 34]. Scaling the distribution of movement displacements generated for different imaging frequencies also allows to determine the type of cell walk because Levy walks-based movement distributions are thought to be relatively invariant to such rescaling [6, 8, 33]. Such methods, however, have not been applied to quantitatively characterize the movement program of T cells in the liver.

While several studies provided strong evidence that T cells undergo a specific movement strategy mechanisms explaining such movement strategy in general remain unknown. Constrained environment of the skin epithelium is likely responsible for the movement patterns for skin-resident CD8 T cells [21]. Activated CD8 T cells in murine brains undergo Levy walks [8], however, whether such a walk type is driven by cell-intrinsic program (as suggested by Harris *et al.* [8]) or arises due to other factors has not been determined. Indeed, previous ecological studies suggested that turbulence of fluids or air may explain patterns of movement of individual cells or birds, respectively, but the actual features of turbulences that allow agents to undergo Levy walks were not directly measured [35, 36]. Furthermore, in a theoretical study Volpe & Volpe [37] also showed that environmental constrains impact the movement patterns of agents and efficiency of the search for targets suggesting that in addition to cell-intrinsic programs other factors are likely to impact movement patterns of T cells [20]. The importance of physiological factors such as blood flow in highly vascularized tissues such as brain or liver in determining movement patterns of T cells as far as we know has not been evaluated.

In this study with the use of intravital microscopy of live mice we provide the first accurate quantification of the movement pattern of liver-localized CD8 T cells (and openly share these data with the community). We show that in most circumstances CD8 T cells undergo short, Brownian-like displacements characterized by a thin-tailed distribution that yet result in transiently superdiffusive displacements, i.e., most T cells in the liver undergo correlated random walks. Importantly, however, the duration of the transient superdiffusion is longer than that observed in other systems (e.g., by T cells in LNs or murine brains). By accurately quantifying the structure of liver sinusoids we show that transiently superdiffusive behavior of liver-localized T cells naturally emerges for Brownian-like, persistent walkers when simulations of movements are done in the liver. Interestingly, our analysis of the structure of fibroblastic reticular cells of LNs suggested that LN-localized T cells should be superdiffusive for similar lengths of time as liver-localized T cells; however, LN-localized T cells are superdiffusive for only few (~ 5) minutes suggesting that other factors such as width of the FRC paths or chemical cues may also reduce the duration of movement persistence of T cells in the LNs. Interestingly we found that in some circumstances T cells in the liver may also undergo Levy flights – large displacements occurring nearly instantaneously – due to de-attaching from the sinusoidal epithelium and flowing with the blood flow. Taken together, we have carefully characterized movement patterns of liver-localized CD8 T cells and provided examples of how environmental constrains (structure of liver sinusoids) and physiology (blood flow) may regulate the observed T cell displacements.

## Materials and Methods

### Experimental Details

#### T cell movement in the liver

Experimental data on movement of wild type (WT), GFP-expressing, TCR transgenic CD8 T cells (OT-I) have been generated similarly as in our previous study [31]. OT-I cells were activated *in vitro* as follows. Splenocytes were incubated with 1 *μ*g/ml SIINFEKL peptide (Biomatik) at 37°C for 30 minutes, washed, and cultured in complete RPMI 1640 (10% FCS, 2mm L-glutamine, 1mm Na-Pyruvate, 100U/ml Penicillin/Streptomicin, 5mm HEPES, 20 *μ*g/ml Gentimicin and 50 *μ*M *β*-mercaptoethanol) at 37° C, 5% CO_2_. Two days later cells were passaged and cultured in fresh media supplemented with (12.5U/ml) recombinant IL-2 (PeproTech). Cells were passaged again after 24 hours, and given fresh media with IL-2. Four days after activation cells were ready for collection and use. Cells were collected from tissue culture flasks, washed, layered over Histopaque^®^ 1083 (Sigma) and centrifuged at 400rcf, for 30 minutes at 21°C to remove dead cells. Live lymphocytes were collected from the interface between the Histopaque^®^ and cell suspension media. Cells were washed a further 3 times prior to transfer to mice.

To label the sinusoids, Evans blue or Rhodamine-dextran were injected i.v. into mice (50 *μ*g/mouse) immediately prior to imaging of the livers (Fig. 1A&B). In some cases, static images of the livers were acquired using two phone microscope. In total, we have generated images of the liver sinusoids from 5 mice (with or without T cells).

**Figure 1:**
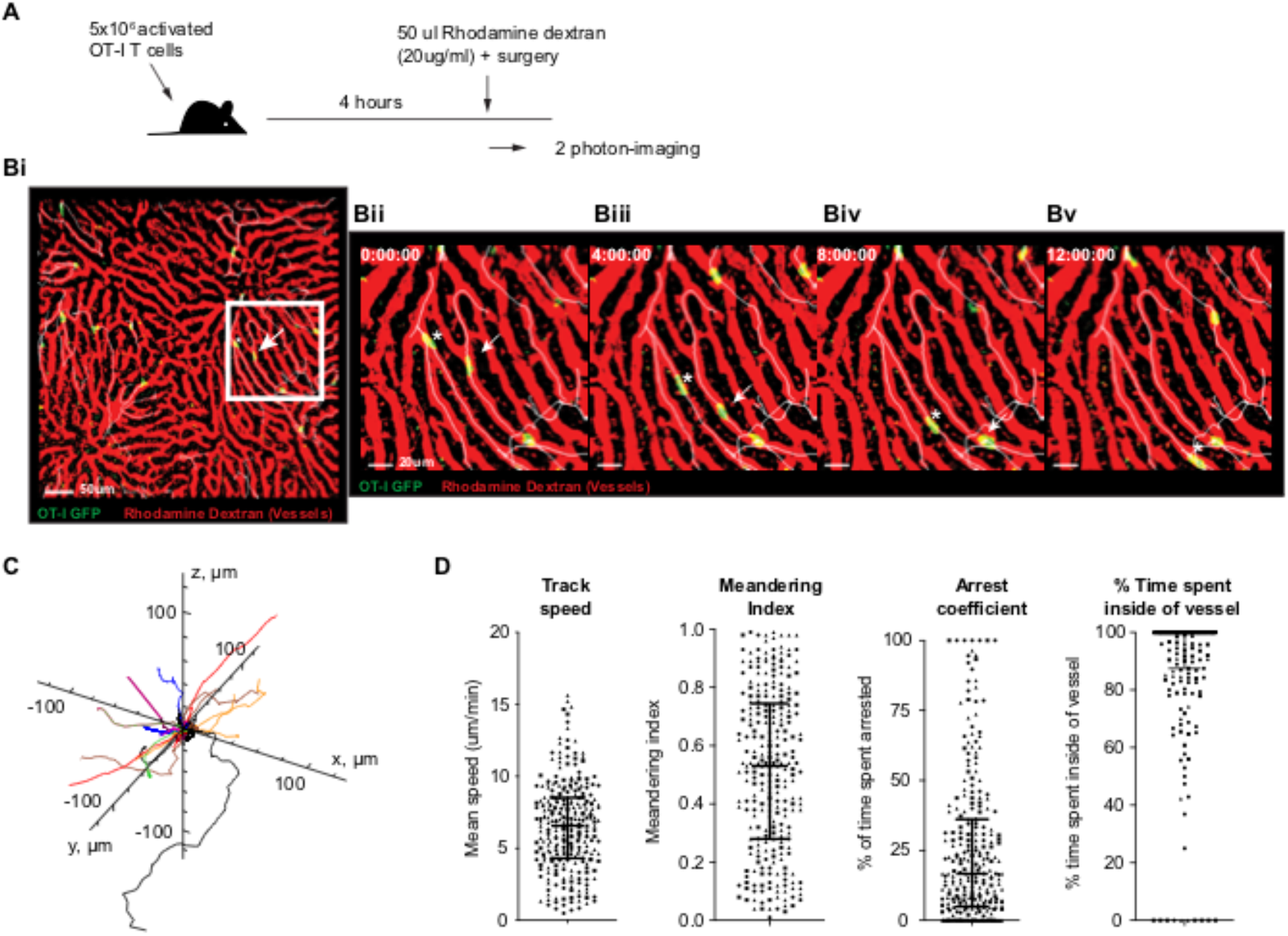
The majority of *in vitro* activated CD8 T cells transferred to naive recipients move in liver sinusoids. We performed intravital imaging of the livers of mice that had previously received *in vitro* activated CD8 T cells and i.v. injection of rhodamine dextran (see Materials and Methods for more detail). (A) 4 hours after i.v. cell transfer of 5 × 10^6^ OT-I GFP CD8 T-lymphocytes (green), mice were injected with 50 μl (20 μg/ml) of rhodamine dextran 70kDa (red) and immediately prepared for intravital imaging. The mice were imaged using a two-photon microscopy with a standard galvanometer scanner in order to acquire a 50 μm deep z-stack every 13 or 26 sec. (B) A sequence of images taken over time to demonstrate the movement of GFP cells (green) through the sinusoids (red) in the direction indicated by the white lines (Movie S1) with star and arrow indicating displacement of two different cells. We show the overall scanned area (Bi) and time lapse for 2 cells (indicated by an arrow and a star) moving in the sinusoids (Bii-Bv). Scale bar is 50 μm (Bi) or 20 μm (Bii-Bv). Time in min:sec:milliseconds is shown. (C) Tracks of randomly chosen 30 cells are shown. (D) Movement parameters of OT-I GFP cells in the liver sinusoids such as average track speed, meandering index, and arrest coefficient (*n* = 269). For every cell movement we calculated the percent of time each cell spent in the sinusoids by calculating signal overlap between cell fluorescence (GFP) and sinusoid fluorescence (rhodamine dextran, *n* = 231). The imaging data in B is representative of 3 experiments, and analysis shown in D is for datasets pooled together (means and SD are shown by bars).

Mice were prepared for multi-photon microscopy essentially as previously described [31]. Briefly, mice were anesthetized with a mix of Ketamine (100 mg/kg) and Xylazine (10 mg/kg). Throughout the surgery and imaging procedure the mouse temperature was maintained at 37°C using a heating mat attached to feedback probe inserted in the mouse rectum. A lateral incision was made over the left lobe of the liver and any vessels cauterized by applying light pressure to the vessel until clotting occurred naturally. The mouse was then placed in a custom made holder. The liver was then exposed and directly adhered to a coverslip that was secured in the holder. Once stable the preparation was transferred to a Fluoview FVMPE-RS multiphoton microscope system (Olympus) equipped with a XLPLN25XWMP2 objective (25x; NA1.05; water immersion; 2mm working distance). For analysis of the motility of liver-localized CD8 T cells a 50 *μ*m z-stack (2 *μ*m/ slice) was typically acquired using a standard galvano-scanner at a frame rate of 2-4 frames per minute.

Other datasets for movement of WT and LFA-1-deficient CD8 T cells, or WT CD8 T cells 1 or 4 weeks after immunization with radiation-attenuated *Plasmodium berghei* sporozoites Pb-CS^5M^ were generated previously [31].

#### Imaging data processing and basic analyses

Raw imaging data was analyzed with Imaris x64 software (Bitplane) v9.12. Tracking of individual cells in a z-stack was performed using the ‘Spots’ function in the surpass mode, and in a single slice by the ‘Surfaces’ function in surpass mode. Detection of individual cells relied upon their relative fluorescence intensity, size (diameter ≥ 9 *μ*m) and used the remaining default function settings for the calculation of the motion tracks using an autoregressive motion algorithm. Tracks were also manually inspected and curated. Basic cell characteristics such as average track speed, meandering index and arrest coefficient were calculated using Imaris. To calculate the percentage of T-cell movements that were occurring in the sinusoids we generated a co-localization channel which calculated colocalized signal of both red and green fluorescence within each frame. To do this, we utilised the colocalization function on Imaris with the threshold defined in Imaris as 16 for uGFP OT-1 cells which subtracted background fluorescence of the liver, and 3000 for rhodamine dextran. We then calculated the total number of frames a T-cell was tracked using the unique cell ID, and compared this to the total frames each individual T-cell was detected in the designed colocalization channel. Rhodamine dextran fluorescence was dependent on the depth of imaging, and in the lower Z-stacks the frequency of T-cells with colocalized signal was poor (results not shown).

#### Data details and data cleaning steps

In our analysis we used 3D coordinates of Plasmodium-specific CD8 T cells in the liver over time from the following experiments:

1. Novel data on movement of GFP-expressing activated CD8 T cells called “New13” and “New26”. In the “New13” dataset data were generated from three mice and frames were sampled at 14.63 (14.63-14.64) sec apart. In the “New26” dataset data were generated from two mice, and frames were sampled on average at 25 (24.29, 25.51) sec apart. To increase the power in the analysis we generated a combined dataset “New1326” by merging data from datasets “New13” and “New26”. We ignored the mid time-points in time-frames in the “New13” dataset, thus, computing time-frames approximately at 26 sec frequency.
2. Previously published data on movement of WT and LFA1-deficient CD8 T cells in the liver [31] resulted in “Lfa1 WT” and “Lfa1 KO” datasets. Data were from 7 mice, and frames were sampled, on average, at 27 (24.65 - 29.56) sec apart.
3. Previously published data on movement of *Plasmodium*-specific CD8 T cells following vaccination of mice with radiation-attenuated sporozoites [31]. In “Wk1” dataset data were generated from two mice immunized with RAS 1 week before. Frames were sampled, on average, at 6.51 (6.51-6.51) sec apart. In “Wk4” dataset five mice were RAS-immunized 4 weeks prior to imaging, and frames were sampled, on average, at 3.25 (3.25-3.26) sec apart.
4. Novel data on movement of *Plasmodium*-specific (OT-1) and lymphocytic choriomeningitis virus-specific (P14) CD8 T cells moving in livers of *Plasmodium berghei*-infected mice [52].

In the Imaris-processed movies we found that there were random missing time-frames in tracks in nearly all data. This could happen when Imaris would associate a cell that went out of focus for one time step with other time steps. Because having a consistent time step to determine cell movement characteristics such as mean squared displacement and movement length distribution was critical we cleaned the data in the following ways. We split tracks with missing time frames and converted them to individual cells for the analyses. For example, if one time frame was missing for a cell, the track was split into two with the second part being assigned as a new “cell”. This created many more cells with shorter tracks. Taken together, cleaning up the data yielded the following. For “New13” dataset we originally had 56 T cells out of which we generated 14 new split cells with 1-2 splits resulting in total of 70 T cells with 2565 time-frames (movements). For “New26” dataset we had 213 T cells with 51 new split cells with 1-2 splits from the original tracks totaling to 264 T cells, with 7736 time-frames. For the combined “New1326” dataset we had 334 T cells with 9018 time-frames (smaller number of time frames in this dataset as compared to New13+New26 is due to different in timestep in two datasets). For “Lfa1 WT” dataset we originally had 198 T cells out of which we generated 19 new split cells with 1-2 splits from the original tracks totaling to 217 T cells, with 6210 time-frames. For “Lfa1 KO” dataset we originally had 154 T cells out of which we generated 16 new split cells with 1-2 splits from the original tracks totaling to 170 T cells, with 5097 time-frames. For “Wk1” dataset we had 99 T cells and generated 3 new split cells with 1-2 splits from the original tracks, totaling to 102 T cells, with 2525 time-frames. For “Wk4” dataset we had 44 T cells with 4169 time-frames that required no cleaning. Most of the analyses have been repeated on “clean” data in which tracks with missing time-frames were removed with similar conclusions reached.

#### T cell movement in the LNs

Data have been provided by Mario Novkovic from their previous publication [46]. In short, 3 × 10^6^ naive P14 TCR transgenic T cells were transferred i.v. into naive mice. Three to twenty-four hours after T cell injection, intravital imaging was performed on the right popliteal LNs. To detect FRCs, anti-YFP mAbs were used and samples were imaged using a confocal microscope (LSM-710; Carl Zeiss)[46]. In total we had 3 sets of images of LNs with FRCs labeled (about 30 images per set with 2 μm depth). In addition, we analyzed 3D movement data of T cells in the LNs (in total 6 movies) produced in the same publication [46]. The data were also cleaned and tracks for which there were not always consecutive timeframes available were split as described above. Analyses were repeated with “clean” data in which cells with missing time-frames were removed; similar results were found. These data were generated at frequency of 20 sec per frame.

### Modeling and Statistical Analytical Methods

#### Mean squared displacement, turning angles, and velocity correlations

For each of the datasets we calculated several basic characteristics. To calculate mean squared displacement (MSD) we used all tracks in the dataset in which time frequency of measurements was approximately the same:

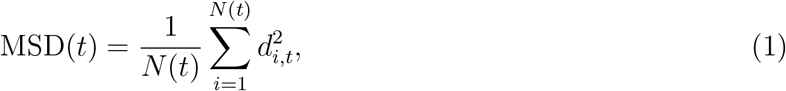

where *d*_*i,t*_ is the displacement of an *i*^*th*^ cell from its initial position to the position at time *t* and *N*(*t*) is the number of cells for which coordinate data were available at time *t*. Note that this is different from movement lengths — which are movements T cells make between sequential time frames. To characterize the MSD we used the following relation MSD(*t*) = *ct*^*γ*^ where *t* are lag-times or delays, and *γ* was estimated by log-log transforming MSD and *t*. Estimate of *γ* < 1 suggests subdiffusion of cells, whose walks are constrained by boundaries, regardless of their step length distribution, *γ* = 1 suggests Brownian (or normal) diffusion, whose walks are affected by neither boundaries nor transportation, whereas, *γ* > 1 suggests superdiffusion, whose walks are indicative of transportation or directed movements. Root mean squared displacement, *ζ* = RMSD was calculated as

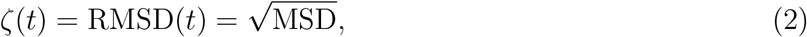

where MSD is defined in eqn. (1). To calculate turning angle for cell movement between time steps (*t, t* + 1) and (*t* + 1, *t* + 2) we calculated the vectors of movements between two time points and then calculated the angle between these two movement vectors. To estimate decay in the correlation movements of a T cell over time we calculated the vector for the initial T cell velocity 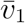 (movement vector for a cell from *t* = 0 to *t* = 1), vector for later movement 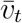 (movement vector for a cell between times *t* − 1 and *t*), and the angle *ϕ*_1,*t*_ between the two vectors 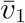 and 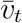:

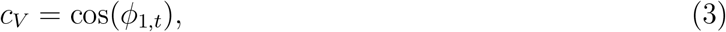

where by definition *c*_*V*_ varies between −1 and 1.

#### Basic statistical distributions to describe movement length data

To investigate which model is best consistent with the distribution of movement/step lengths of activated CD8 T cells in the liver or distribution of branch lengths of liver sinusoids we fit several alternative distributions to the data. Cauchy distribution is defined as

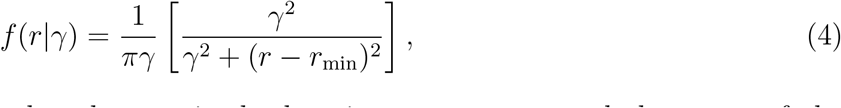

where *r* ≥ *r*_min_ is the movement length, *r*_min_ is the location parameters, and the mean of the distribution is undefined. Half-gaussian distribution with *r* ≥ 0 is defined as

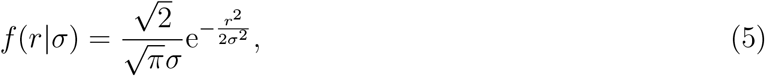

where mean of the half-normal distribution is 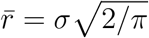.

Stable Levy (SL) distribution is defined as

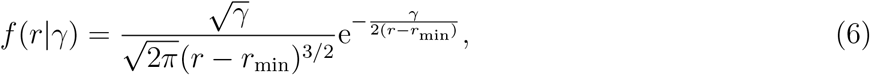

where *γ* and *r*_min_ are the shape parameter and location parameter, respectively. The SL distribution has infinite mean and variance.

Generalized Pareto (GP) distribution is defined as

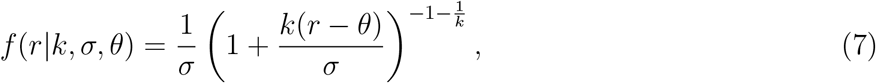

where *k* is the shape parameter, *σ* is the scale parameter, and *θ* is the location parameter. In this distribution, the mean is 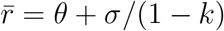.

When *k* = 0 and *θ* = 0, GP becomes exponential distribution

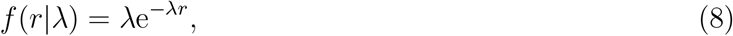

where *λ* = 1/*σ*, and mean being equal to 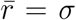. When *k* > 0, *θ* → 0 GP becomes the Pareto (or Powerlaw) distribution

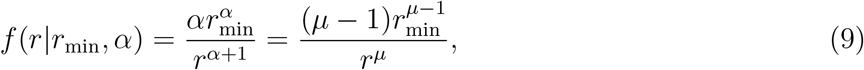

where *r*_min_ = *σ/k* (scale parameter), *α* = 1/*k*, *μ* = *α* + 1 (shape parameter), and 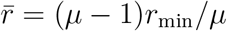. Note that in Pareto distribution *r ≥ r*_min_. The important property of the Pareto distribution, describing either Brownian or Levy walks, is dependence of the mean and variance on the value of *μ* (and thus *α*). For *μ* > 3 (*α* > 2), both mean and variance of the Pareto distribution are finite corresponding to Brownian walk. In contrast, when *μ* ≤ 3 (*α* ≤ 2) mean is finite and variance is infinite corresponding to Levy walks. Finally, when *μ* ≤ 2 (*α* ≤ 1), both mean and variance are infinite corresponding to bullet motion [1].

Generalized extreme value (GeV) distribution is defined as

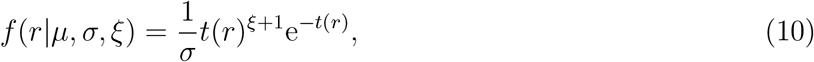

where 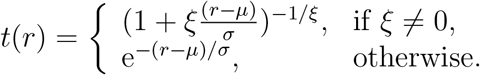

#### Statistical methodology of fitting models to data

We fit the models (eqns. (4)–(9)) to either actual movement lengths *r* of cells between two time points or scaled movements *ρ*. Scaled movements are defined as 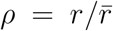, where 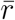 is the average movement in the particular dataset (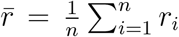 for *n* movements). Using scaling of movement lengths by square root of mean of squared movement lengths 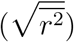 gave similar results. It is important to emphasize that scaled movements *ρ* allows to convert cell tracks to be independent of the effect of different mean speeds of cells and independent of the effect of different sampling frequency [8]. Likelihood of the model parameters given the data is given as

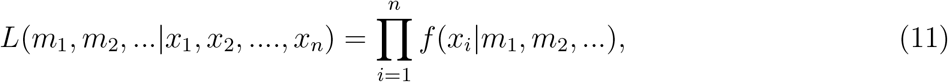

where *x*_*i*_ are either the cell movement length data *x*_*i*_ = *r*_*i*_ or the scaled movement data *x*_*i*_ = *ρ*_*i*_(*t*) = *r*_*i*_/*ζ*_*i*_(*t*), for *i* = 1*...n*, for *n* data points, for fixed time intervals *t*. The probability density functions *f*(*x*_*i*_|*m*_1_, *m*_2_, ...) are given in eqns. (4)–(9). Parameters *m*_1_, *m*_2_, ... of the models were estimated by minimizing the log of negative likelihood, 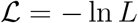.

#### Tail analysis of the step length distribution

In addition to fitting different distributions to the movement length/displacement data we performed an analysis to determine the shape parameter of the powerlaw/Pareto distribution *μ* that is best consistent with tail of the displacement distribution data [34]. Given that Pareto distribution has two parameters (*r*_min_ and *μ*, eqn. (9)) one needs to define *r*_min_ that determine the cut-off in the data and that allows for the best description of the tail of the data with Pareto distribution. The steps to define *r*_min_ are as follows [34]: (1) for each possible choice of *r*_min_ from the smallest to the largest unique values of *r*, we computed *μ* by maximum likelihood estimate given by the analytic expression 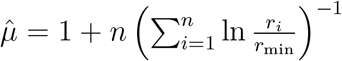, where *r*_*i*_, *i* = 1 ... *n*, are the observed values of *r* and *r*_*i*_ ≥ *r*_min_. In doing so, we also set a limit to the number of data points in the analysis by setting a criteria by which the analysis is terminated if the standard error on 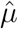, approximately given by 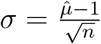, was greater than 0.1, as higher order corrections of the standard error are generally positive. This condition was set because the estimates of *μ* can be erroneously large when *r*_min_ is so large that the number of observed *r* ≥ *r*_min_ is very small. It was also assumed that *μ* > 1, since the distributions with *μ* ≤ 1 are not normalizable, and hence not realistic in nature. We selected *r*_min_, at which the Kolmogorov-Smirnov (K-S) goodness-of-fit statistic *D* was the minimum out of all *D* values given by the fits over all possible unique *r*_min_’s of the data. Here, we also made a finite correction to the estimates of *μ* for small sample sizes (*n* < 50) in addition to the condition set by the standard error of *μ*.

From the above analyses, we carried forward the estimated *r*_min_, *μ*, and *D* to the next stage of testing the probability of the data, given that the estimated model is the plausible one, by the method of Kolmogorov-Smirnov goodness-of-fit test. To do that we generated data from the hypothesized (or estimated) Pareto model, sampling using a semi-parametric bootstrap method, computing *D* for each sample, which is the synthetic *D*, and thus, computing a fraction of those were greater than the empirical *D* of the given data, which yields the *p* value of the hypothesis. We then computed the uncertainty in the estimates of *r*_min_, *μ*, and *n*, given by standard deviations, by resampling the data with replacements by Markov chain Monte Carlo (MCMC) simulations.

#### Digitizing and skeletonizing the 3D sinusoidal structure

We analyzed 23 z-stacks of Evans blue stained 512 × 512 *μ*m images of the liver (mouse 1: 171207 and mouse 2: 190624), as well as 3 sets of images/movies with T cells in which liver sinusoids were labeled with rhodamine-dextran. Resulting structures were similar based on calculated characteristics such as branching angle distribution and straightness index (results not shown). Light-adjusted 2D images were converted from RGB into grayscale using rgb2gray, filtered through Matlab filters (1) imfilter with Gaussian N(10,2) to remove noise, (2) medfilt2, with ‘symmetric’ setting to remove all the lines from the image, (3) uint8 to subtract image without lines from image with lines, (4) setting threshold at *>* 130 of the subtracted image, and (5) removing more noise by imopen, and saving all 2D images in a stack converting to a digitized 3D image, with “1” giving the sinusoids and “0” giving the hepatocytes and other unstained tissues (Fig. 3). The digitized 3D image was then skeletonized with 1’s giving the mean path of sinusoids and 0’s elsewhere, converting to a voxel image, using parallel medial axis thinning method as per the MATLAB vectorized implementation of the algorithm [59]. The 3D binary voxel skeleton was converted to a network graph, computing nodes (junctions) and edges (branches), using previously published methods [60], and mapping the respective skeleton to 3D graphics function. The nodes and the branches were used to compute branch angles, branch lengths, node-to-node direct distances, number of branches per node. Straightness of branches metric was computed as the node-to-node direct distance over the branch length. The binary 3D liver image was used to run simulations of cells walking on the 1’s on the lattice, scale of which can be decided, for e.g., 1 unit equals 0.01 μm (Fig. 4).

#### Simulation methods of cell walks on digitized sinusoidal structures and analyses of re-sampled step-length distribution and MSD

We simulated cell walks on (i) digitized 3D real sinusoidal branch structures (5 imaging sets) and (ii) no structures (5 simulations) imposing no physical constraints giving freedom for cells to turn any direction after every step. For each generated structure we selected a random point on a mean sinusoidal path, i.e. on the digital skeleton given by 1’s, on the matrices generated from digitized 3D sinusoidal structures of 1’s and 0’s in the real case, and on a branch position coordinate in the simulated case, as the starting point of a T cell. Then we selected a cell step-length from the Pareto distribution with different parameter *μ*. Note that we considered all movements as forward movements, as reverse movements of cells along the sinusoids were the minority in the data (about 20%, Fig. S3). Ability of cells to move in one preferred direction in constant external conditions can be viewed similarly to the physics postulate of momentum conservation (or “biological inertia”, [61]). We disregarded deviation of tracking cell centers from the mean sinusoidal path because the cell diameter, on average, 10 *μ*m, was greater than the cylindrical diameter of the sinusoids, 7 *μ*m, and thus, T cells likely occupied most of the cross-sectional area of the sinusoids. Note also that step-lengths and branch lengths were thus simulated disregarding their autocorrelations assuming they are independent and identically distributed (i.d.d.). Cell walk simulations on no-structure, the cell turning angles were random from a uniformly spherical distribution. Note that the walk simulations of cells and sampling coordinate were terminated once a cell hits the boundary of the structure before the simulation time was elapsed in the case of real sinusoidal structures. We computed MSD of all simulation scenarios. To simulate T cell walks in digitized 3D network of FDCs in LNs we proceeded similarly to the method for liver sinusoids assuming that T cells can only walk in the middle of the network path and turns can be taken only when a T cell reaches a branching point.

#### Simulation methods of computing efficiency of cells finding a target

To calculate efficiency of the T cell search for the malaria liver stage we calculate the time-to-target as this is being most relevant for CD8 T cell-mediated protection against malaria. In simulations, T cell started search from a random point but with a given distance to the parasite. We assumed that T cell found the infected cell when it was within 40 *μ*m from the target, an approximate size of the murine hepatocyte [53, 54]. We repeated the simulation with 5 different distances to targets (results not shown). For 3D simulations if searching T cell reached the boundary we assumed that it missed the target (and thus was not included in the calculation of the average time). For simulating T cells searching for a target in the liver we simulated T cells moving forward on one path, effectively reducing search problem to 1D. This is reasonable given that difference between Brownian and Levy walks is only on the movement length distributions and not on turning angles that cells take [1]. To simulate *n* T cells searching for a single sample, assuming that search is done by different cells independently, we resampled times to target with replacement *n* times from the dataset on a single T cell searching for a single target, and calculated minimal of the time in the sample. This procedure was them repeated 1,000 times to generate a distribution to target when *n* T cells search for the infection site.

## Results

### Majority of liver-localized CD8 T cells crawl in liver sinusoids

To understand what strategies *Plasmodium*-specific CD8 T cells may be using to survey the liver for infection we have established a protocol to generate stable numbers of liver-localized CD8 T cells [31]. In these experiments, naive CD8 T cells are activated *in vitro* and then transferred i.v. into naive mice that either left uninfected or can be later infected with *Plasmodium* sporozoites (Fig. 1A) [31]. By injecting fluorescent label (rhodamine dextran) i.v. immediately prior to surgery and imaging we can efficiently label the liver sinusoids (Fig. 1B). Intravital imaging showed that CD8 T cells in the liver are highly motile cells with the average speed of 6.6 *μ*m/min, a high meandering index of 0.52, and a low arrest coefficient of 0.25 (Fig. 1C and Movie S1). Importantly, these CD8 T cells were observed to be located inside the liver sinusoids most of the time (89-100%, Fig. 1D). Due to incomplete labeling of the sinusoids by rhodamine dexran and reduced red signal from deeper liver areas it is likely that a lower bound is an underestimate and in this system nearly all T cell movements (and by inference, most T cells) were localized in the sinusoids and are, thus, intravascular. As compared with the blood flow rate in the liver, a relatively low average speed of these liver-localized T cells and T cells in our previous experiments with high frequency intravital imaging suggests that the cells are crawling on the endothelial wall of the sinusoids and not flowing with the blood [31]. Thus, our robust experimental set-up allows for the first time to accurately quantify movement patterns of liver-localized CD8 T cells along with the well defined structures (i.e., hepatic sinusoids).

### Most liver-localized CD8 T cells undergo correlated random walks

In our experiments movements of liver-localized *Plasmodium*-specific CD8 T cells were recorded with two photon microscopy. Interestingly, T cells in the liver displayed transiently superdiffusive overall displacement characterized by MSD increasing faster than linearly with time (with slope *γ* = 1.55) for up to 20 min (Fig. 2A). Saturation in MSD with time in these and other experiments typically arises in part due to rapidly moving T cells leaving the imaging area (Fig. S1). Superdiffusion is a hallmark feature of Levy walks [3, 38], and yet liver-localized T cells only displayed short, Brownian-like movements (Fig. 2B). Tail analysis [34] of the distribution of actual or scaled movement lengths (movement lengths normalized by mean movement length) suggested the shape parameter of the powerlaw distribution *μ* = 5.6 — an indication of Brownian (*μ* > 3) and not Levy (*μ* ≤ 3) walks. Interestingly, scaling the movement length distribution for sub-sampled data (i.e., for data sampled at a lower frequency than was used at imaging) resulted in relatively similar distributions (Fig. S2) suggesting that even thin-tailed movement length distributions may be similar for sub-sampled, scaled data. Thus, large overall cell displacements with small movement lengths arose not due to Levy-like cell-intrinsic behavior but due to small turning angles and a strong correlation between initial and subsequent velocity vectors for at least 10-15 min, resulting in cells moving in one direction for substantial periods of time (Fig. S3).

**Figure 2:**
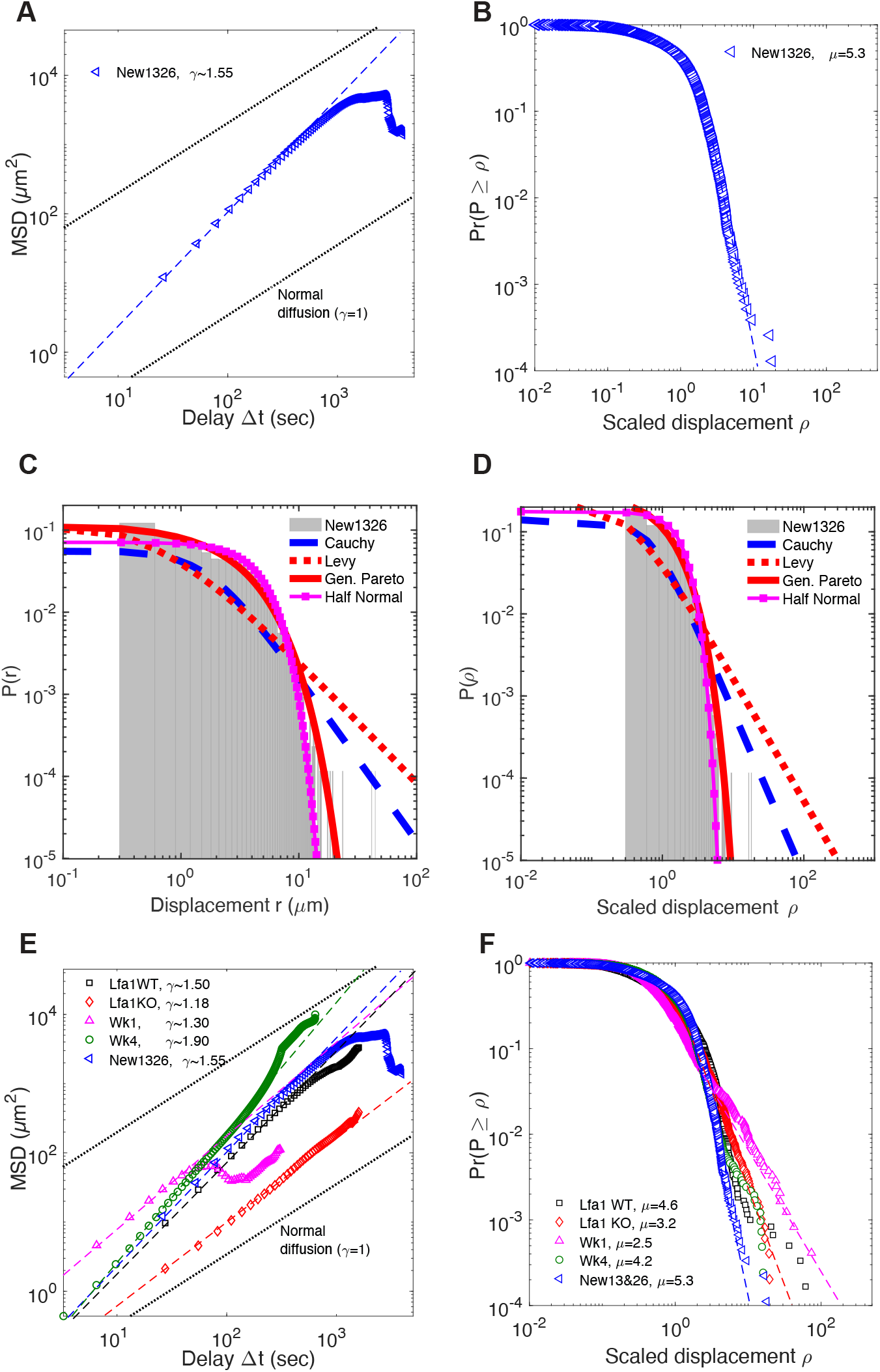
Activated CD8 T cells in the liver are Brownian walkers displaying super-diffusive displacement. For experiments described in Fig. 1 we calculated basic movement characteristics such as the mean squared displacement (MSD, A) and distribution of scaled movement lengths (B) for all T cells (*n* = 268). Resulting slopes for linear regressions for the log(MSD) with log(*t*) and movement length distribution (*γ* and *μ*, respectively) are shown on individual panels. (C)-(D). We fit several alternative models of movement length distribution to the raw (C) or scaled (D) displacement data from experiments in Fig. 1 (eqns. (4)–(9)). Parameters for the fitted models along with AIC values are given in Tabs. S1 and S2. (E)-(F). We performed similar analyses with other published data for *in vitro* or *in vivo* activated CD8 T cells in the liver [31]. Cell displacement in the liver was characterized by MSD with slope *γ* (E) and movement lengths characterized by the tail slope *μ* (F). Additional details on analysis of movement length distribution are given in Fig. S4. Scaling in (B) and (F) was done for all tracks per dataset using mean displacement length 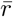 using relationship 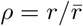 (see Materials and Methods for detail).

To complement the movement length distribution tail analysis – which is the standard in the field [7, 8, 12] – we fitted a series of alternative mathematical models characterizing the distribution of movement lengths of T cells. We found that generalized Pareto distribution fitted the data with best quality (Fig. 2C-D and Tab. S2). Best fit parameters of the generalized Pareto distribution predicted a finite mean and variance for the movement length distribution – which again are the chief features of the Brownian and not Levy walkers [1]. Thus, in our experiments liver-localized CD8 T cells performed correlated (persistent) random walks with transient superdiffusive movements that were longer than those observed for CD8 T cells in the brain [8].

To verify that this result was not due to a potential impact of the injected rhodamine-dextran dye we analyzed four other datasets from our previous study on movement of liver-localized CD8 T cells [31]. These experiments involved transfer of *in vitro* generated wild-type and LFA-1-deficient activated CD8 T cells or *in vivo* generation of liver-localized T cells by transfer of naive OT-I cells to mice followed by immunization of these mice with radiation-attenuated malaria sporozoites (RAS) expressing the SIINFEKL epitope recognized by the T cells [29, 31]. Both *in vitro* wild-type activated cells and *in vivo* activated cells imaged at a memory time point (4 weeks) after immunization also displayed transiently superdiffusive behavior with *γ* > 1 for 10-20 min (Fig. 2E). Interestingly, LFA1-deficient T cells which cannot strongly attach to endothelial cells and roll under the blood pressure [31], displayed displacement closest to Brownian and yet were transiently superdiffusive (*γ* = 1.18 in Fig. 2E). We also observed high heterogeneity in MSD in mice imaged one week after RAS immunization (Fig. 2E). At this time point after *in vivo* priming, many CD8 T cells do not undertake the crawling behavior characteristic of liver-localized T cells, most likely due to intermediate expression of LFA-1 shortly after immunization [31].

Likewise all but one cell population had relatively small movement lengths. The exception was for CD8 T cells in mice one week post RAS immunization (Fig. 2F) where cells do not attach firmly to the liver endothelium and often flow in the bloodstream, thus, performing Levy flights (*μ* < 3, Fig. 2F and [31]). To detect such floating events we imaged the livers at higher than usual frequency (3-6 frames/sec). Removing displacements that occur at high speeds (e.g., ≥ 40 *μ*m/min resulted in movement length distribution being consistent with thin-tailed distribution (i.e., Brownian-like movements) and yet transiently superdiffusive MSD (results not shown). Transiently superdiffusive behavior with small, Brownian-like movements was also observed for *Plasmodium*-specific activated CD8 T cells and activated CD8 T cells with irrelevant specificity in *Plasmodium berghei*-infected mice (Fig. S5). Taken together, our results establish that most crawling liver-localized CD8 T cells display correlated random walks resulting in transiently superdiffusive behavior and small movements [39]. T cells, that poorly attach to the sinusoidal endothelium undergo a combination of correlated random walk with Levy flights also resulting in superdiffusion and fat-tailed distribution of movement lengths.

### Liver sinusoids have short lengths and small branching angles

Given the network structure of liver sinusoids (Fig. 1B) and universal pattern of transient but relatively long superdiffusive behavior of liver-localized CD8 T cells we hypothesized that a linear structure of the sinusoids may be forcing T cells to exhibit persistent walks. Therefore, we analyzed 3D structure of liver sinusoids. For these intravital imaging experiments we injected a contrast dye (Evans blue or rhodamine-dextran) and imaged murine livers (Fig. 3A). Z-stacks of the images were processed in Matlab to generate highlighted sinusoidal maps (Fig. 3B). A subset of the full image was further processed to generate nodes and edges of the sample (Fig. 3C-D). Analysis of the processed digital structures suggested that the branch length distribution was best described by the half-normal distribution with generalized Pareto distribution being second best (Fig. 3E and Tabs. S1 and S2). The longest branch detected did not exceed 100 *μ*m, and the tail of the branch length distribution was characterized by a relatively high shape parameter of the powerlaw distribution (*μ* = 7.65, Fig. 3F) [34]. Importantly, sinusoids tend to branch at a small angle which is likely to improve blood flow (Fig. 3G) [40]. Finally, analysis of the neighboring nodes in the network (Fig. 3D) suggests that sinusoids connecting two adjacent nodes tend to be straight (Fig. 3H). Taken together, properties of the liver sinusoids such as small branching angles, straightness, and shortness appear to match the observed pattern of movement of liver-localized T cells (e.g., Fig. 2F and Fig. 3F).

**Figure 3:**
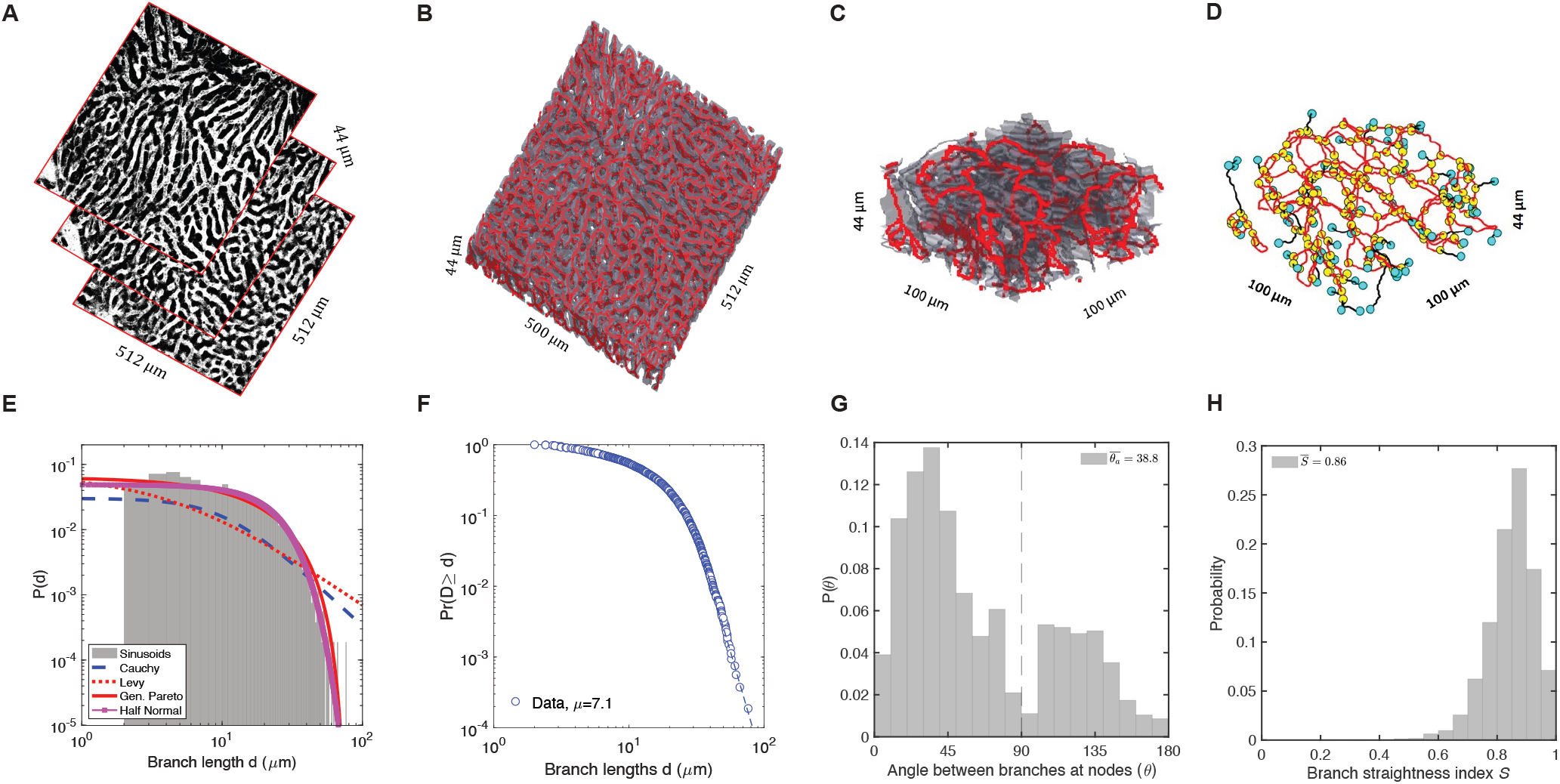
Liver sinusoids have relatively short lengths and small branching angles. We performed experiments in which liver sinusoids were labeled by i.v. injection of Evans blue or rhodamine dextran and then imaged immediately by using intravital microscopy. (A) Z-stacks of 2D images of the liver used for analysis. (B) Computer-digitized 3D sinusoidal maps were generated by contrast thresholding and by stacking up the 2D images. Red shows the center paths of the sinusoids. (C)-(D) Computer-assisted rendering of liver sinusoids for a subset of the data (C, shown in red) used to determine the nodes (D, yellow circles) and the edges/branches (red) of the structure (turquoise circles are the end points). (E)-(H) Characterization of the liver sinusoids. (E) We fit several alternative models to the distribution of lengths of sinusoids using likelihood approach (see Materials and Methods and Tab. S1). (F) Cumulative distribution of the sinusoid lengths was also analyzed using tail analysis with the indicated linear regression slope *μ* [34]. (G) Distribution of branching angles between sinusoids calculated for nodes (*n* = 4013) in D. Parameter 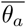 is the average branching angle for acute angles. (H) Straightness of the liver sinusoids (computed as the ratio of point-to-point branch length to actual branch length for all near neighbors in D); 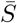 is the average straightness.

### Liver sinusoids alone allow for transient superdiffusion of Brownian-like walkers

To investigate how the structure of the liver sinusoids may impact the movement patterns of cells with an intrinsic movement program we simulated movement of cells assuming that they are either Brownian or Levy walkers (Fig. 4). Specifically, we assumed that the distribution of movement lengths follows a powerlaw distribution with the shape parameter *μ* being different between Brownian (*μ* = 4.5) and Levy (*μ* = 2.5) walkers while fixing the average movement length for two walk types which is possible when *μ* > 2 [1]. As expected in free 3D space Brownian walkers displayed normal diffusion (*γ* = 1) while Levy walkers superdiffused (*γ* = 1.5, Fig. 4A-C). In contrast, when cells were run on the digitized network of liver sinusoids (Fig. 3A-D) with cells allowed to make a random choice of turning when reaching a branching point the movement pattern changed dramatically (compared Fig. 4A vs. D and B vs. E). Both Brownian and Levy walkers displayed superdiffusion in the liver structure for substantial period of time (~ 55 time steps, Fig. 4F) with the diffusion becoming normal at longer time scales. This pattern is similar to small particles displaying transient superdiffusion early and normal diffusion later due to motion of self-propelling bacteria in media [41]. Such a long transient superdiffusion was not due to the assumption of forward movement by Brownian-like walkers because simulations in free 3D space assuming that Brownian-like walkers can only move forward resulted in superdiffusion for only about 7 time steps (results not shown). Thus, the structure of the liver sinusoids along with the requirement for T cells to move forward was sufficient to allow for transient yet long-term superdiffusion of Brownian-like walkers such as crawling liver-localized CD8 T cells.

**Figure 4:**
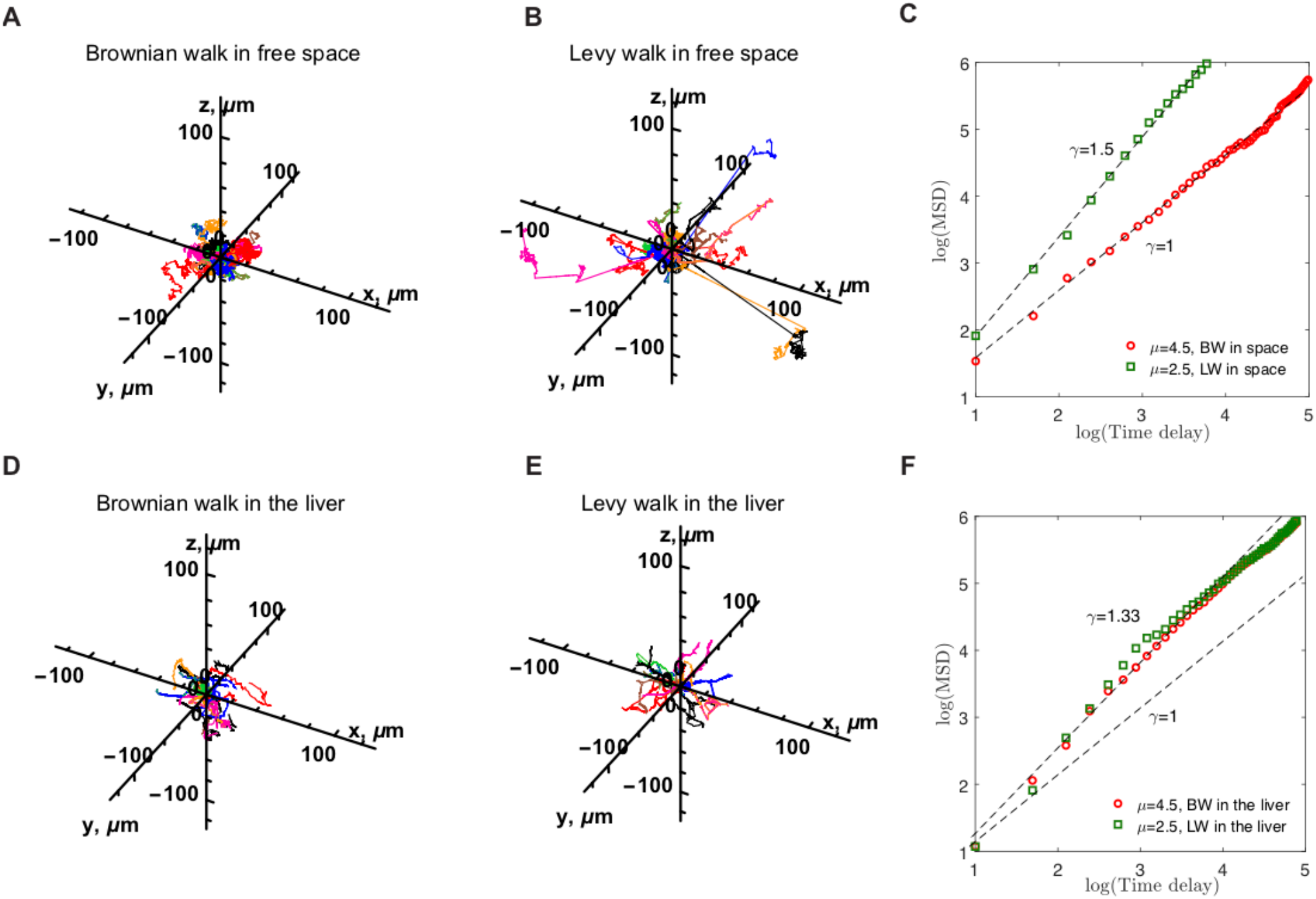
Liver structure imposes strong constraints on the movement pattern and overall displacement for Brownian or Levy walkers. We simulated movement of 200 T cells assuming that their movement length distribution is described by Pareto (powerlaw) distribution (see eqn. (9)) with *μ* = 4.5 (i.e., Brownian walkers, A&D) or *μ* = 2.5 (i.e., Levy walkers, B&E). T cells moved in free environment with no constraints (A-C) or are constrained by liver sinusoids (D-F). A sample of 20 cell trajectories is shown in A-B & D-E. (C)&(F). We calculated the MSD for the walkers and estimated the displacement slope *γ* by linear regression of the initially linear part of the ln-transformed data. In free space simulations turning angle distribution for moving T cells was generated using uniform spherical distribution, while in liver sinusoids movement was set forward along the sinusoidal mean path. The average movement length per time step was 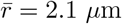 as in our data with time step of 26 sec. An example of a simulated T cell movement in the liver is shown in Fig. S6 (see Materials and Methods for more details).

By finding that the structure of liver sinusoids may be sufficient to allow for transient superdiffusion of T cells we wondered if the same phenomenon may apply to T cells in other tissues. Because previous studies have suggested that T cells in LNs are close to Brownian walkers [3, 6, 42, 43] we sought to determine if this is because of the structural constraints of the LN environment that does not allow for superdiffusion. It has been suggested that T cells use the network of fibroblastic reticular cells (FRCs)/DCs as a guide for movement [44, 45]. Recently, the FRC network of the LNs has been imaged using confocal microscopy [46]. Using a similar computational pipeline we digitized the FRC network using previously published data [46] and calculated its basic characteristics (Fig. S7A-C). Interestingly, similarly to the liver sinusoids the FRC network had relatively short branch lengths with the tail of the distribution being well described by the powerlaw distribution with a shape parameter *μ* = 6.7 (Fig. S7D). A half-normal distribution fitted the data on branch length distribution with best quality (Fig. S7E and Tabs. S1 and S2). Similarly to the liver sinusoids FRC network has relatively small branching angles at the nodes, and connections between adjacent nodes are mostly straight (Fig. S7F-G). By assuming that T cells can only walk in the middle of the digitized FRC networks with preferred movement forward simulations of T cell walks suggested that independently whether cells do Brownian-(*μ* = 4.5) or Levy-like (*μ* = 2.5) walks they display transient superdiffusion similar to that of T cells simulated to walk in the liver (Fig. S7H). This is perhaps unsurprising given how similar the derived characteristics of FRC network is to the liver sinusoids. In contrast, analysis of the experimental data on movement of naive CD8 T cells in murine LNs [46] suggested that such T cells are superdiffusive for very short time period (Fig. S8). The duration of a persistent walk of CD8 T cells in LNs is around 4-5 min while CD8 T cells in the liver superdiffuse for nearly 20 min (Fig. S9). Analysis of the actual moving tracks of T cells in LNs suggests that many T cells tend to turn after a short directed movement (Fig. S8A&E). Indeed, it is well known that naive T cells in LNs tend to be restricted to the T cell areas, most likely by the gradient of chemokines. Also, it is possible that FRC/DC-based scaffolds are wide, thus, allowing for more freedom of T cells to turn. In contrast, liver-localized T cells are likely to be more constrained by the size of the sinusoids, and in the absence of any specific liver infection there are likely not be any chemoattractant cues that would force T cells to turn more. Our results, thus, suggest that while structural constraints of the liver sinusoids or FRC network in LNs may allow for long-term superdiffusion of T cells other factors such as details of the physical scaffolds used for crawling and/or attraction cues in the tissues may impair cell’s ability to superdiffuse.

### Brownian and Levy walkers have a similar search efficiency for a malaria liver stage

Determining the movement strategy of T cells in tissues may be important because different walk strategies may have different efficacies [8–11, 13–17, 33, 37, 43, 47–51]. Search efficiency may depend on details of a specific biological system, however. *Plasmodium*-specific liver-localized CD8 T cells need to locate and eliminate few Plasmodium-infected hepatocytes before these parasites mature and escape into the circulation. We therefore simulated a search by a single liver-localized CD8 T cell for a single *Plasmodium* liver stage assuming that the parasite induces no attraction to the infection site and restricting the average movement speed to be the same for Brownian or Levy walkers. We used Pareto (powerlaw) distribution to describe the movement lengths with scale parameter *μ* = 4.5 for Brownian walk and *μ* = 2.5 for Levy walk (see Materials and Methods for more detail). When a single CD8 T cell searches for the parasite in constraint-free 3D space, on average Levy walkers require less time 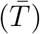 to locate the target (Fig. 5A&C) which is consistent with previous results [8, 9]. Levy walks also result in smaller variability (characterized by the standard deviation of the time to target *σ*) in the time to find the target. Yet, because of the large variability in the time to target, the difference in the time to target between Brownian and Levy walkers in these simulations may seem biologically nonsignificant.

**Figure 5:**
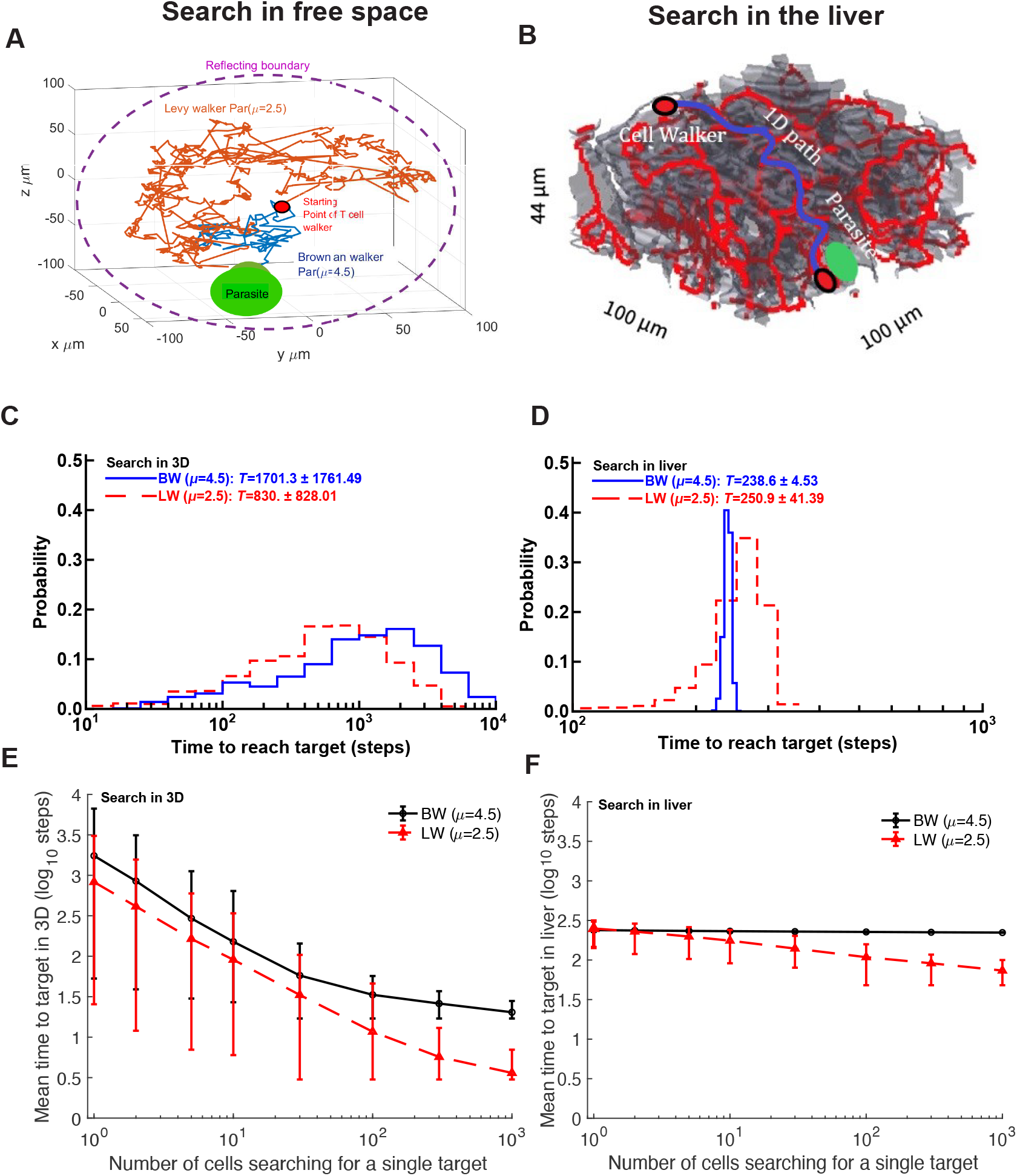
Brownian and Levy walkers have similar efficiency in search for a single malaria liver stage. We simulated search of a T cell for a single target assuming Brownian (BW, *μ* = 4.5) or Levy (LW, *μ* = 2.5) walks in free space (A&C) or in liver sinusoids (B&D) and calculated the time it took for the T cell to locate the target (C&D). We assume that search occurs randomly (no attraction of the T cell to the target). (A)&(C) Levy walkers are on average more efficient in free space. We show distribution of times to the target for simulations when T cell started the search 100 *μ*m away from the target. (B)&(D) Brownian walkers are on average more efficient in the liver. We show distribution of times to the target for simulations when T cell started the search 500 *μ*m away from the target (mean 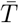 and standard deviation ±*σ* of the time is shown in C&D). (E)&(F) Levy walkers are more efficient when multiple T cells search for a single target. We extended the simulations by allowing *n* T cells to independently search for the infection site. The time to target was calculated as the minimal time to find the target by any T cell out of *n* cells. In all simulations T cell was assumed to find the target when it was within 40 *μ*m from the parasite (an approximate size of the murine hepatocyte). We simulated 1000 cells (3D) or 3000 cells (liver) under each scenario with average movement length per time step being 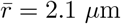 in Pareto (powerlaw) distribution with the shape parameter *μ* (eqn. (9)).

Interestingly, when the search is performed in the digitized liver structures, Brownian-like walkers are on average more efficient at finding targets than Levy walkers (Fig. 5B&D) with smaller variance in the time to target. Furthermore, in 55% of simulations Levy walkers would take longer than the slowest Brownian walker to find the target (Fig. 5D). This may be particularly important in the case of malaria where a failure to find all the targets in the liver within a fixed time period (e.g., 48 hours) will result in blood stage infection and fulminant disease. Alternatively, in the liver Brownian walkers may be unable to locate the infection within a given time period if they start their search too far away from the parasite, while Levy walkers have a non-zero chance (Fig. 5D).

The difference in search efficiency between Levy and Brownian walkers in free 3D space and in the liver is due to differences in the paths to the target. In the digitized liver there are essentially few short pathways between a T cell and the target, and Brownian walkers make consistently small movements moving closer to the target. In contrast, due to small average movement length Levy walkers spend long periods of time pausing before performing a long run sometimes resulting in them reaching the target rapidly. However, long pauses also result in much longer overall time to the target, which increases the average search time substantially.

In these simulations so far we assumed that a single T cell searches for the single infection site. However, in reality multiple T cells may be searching for the same parasite. Indeed, our recent analysis suggests that for 10^5^ liver-localized CD8 T cells in a mouse at least 1 T cell should be within 100 *μ*m of the parasite with 99% certainty [52]. In our experiments we rarely see parasites surrounded by more than 5-8 T cells suggesting that in many situations fewer than 10 T cells are close enough to the parasite to find it within several hours [53, 54]. As expected the average time to find the target declines with an increasing number of searchers, independently whether the search is done in free 3D space or constrained liver environment (Fig. 5E-F). Intuitively, the reduction in time to find the target arises due to variability in search time since we assume that search by individual T cells is done independently; therefore, with many T cells searching for infection, the one T cell having shortest search time will define success of the search and Levy walk allows for a larger fraction of faster searches. Yet, for physiologically relevant numbers of T cells searching for the infection site in the liver (≤ 10/parasite), we find a relatively small difference in the search efficiency between Brownian and Levy walkers (Fig. 5E-F).

## Discussion

T cells utilize various movement strategies depending on the tissue which they are in. By combining novel experiments, data analysis, and computational modeling we have shown that T cells in the liver undergo two types of walk: correlated random walk (by T cells that crawl on sinusoidal endothelium) and a combination of correlated random walk and Levy flights (by T cells, e.g., 1 week post RAS immunization, that poorly attach to sinusoidal endothelium, crawl or float with the blood flow, Fig. 2F). Interestingly, even crawling T cells display transient, yet relatively long-term superdiffusion which we attribute to the special properties of liver sinusoids which constrain movement of liver-localized, intravascular CD8 T cells (Fig. 4). Our results, thus, suggest that observed patterns of superdiffusion or rare long range displacements observed in some experimental data need not arise from cell-intrinsic movement programs but can be explained by the physical and/or physiological properties of the environment in which T cells move (e.g., by floating with the blood flow). Interestingly, while we found that LFA-1-deficient liver-localized CD8 T cells do not firmly attach to sinusoidal endothelium and often flow with the blood flow [31], we did not detect Levy flights for such data (*μ* > 3 in Fig. 2F). This was due to a lower frequency of imaging in these experiments as compared to imaging frequency in experiments week 1 post RAS vaccination (27 sec/frame vs. 6.5 sec/frame, respectively, see Materials and Methods for more detail). Therefore, frequency at which imaging is performed can influence the type of movements cells exhibit.

Specifics of the physical structures in tissues and other factors such as chemoattractant cues are likely to contribute to the movement patterns of T cells in tissues. In particular, we showed that FRC network of the LNs also allows for transient superdiffusion of agents in simulations; however, actual T cells in LNs superdiffuse for very short time periods (Figs. S8 and S9). Thus, other factors such as chemical cues (chemokines), finer details of the physical paths on which T cells walk such as FRCs/DCs, infection-induced inflammation, and preference for forward movement in the absence of external disturbances are likely to be also important in determining overall patterns of T movement in tissues. In particular, the role of environmental cues in determining agent movement patterns has been suggested in several systems [3, 22, 55].

Our results have broad implications for other systems in which superdiffusion and rare long range displacements have been observed or suspected [11, 56]. It will be important to re-examine the evidence supporting intrinsic search programs by matching movement patterns of agents to the experimentally measured environmental details (e.g., airflows for albatrosses or currents for marine predators) and evaluating the role of cues (chemical or otherwise) in defining overall movement patterns of agents [13, 14, 37]. Ultimately, future studies will need to quantify the relative contribution of intrinsic programs, environmental physical constraints, and cues in determining movement pattern of agents [20].

Given our results it is tempting to speculate about potential mechanisms resulting in Levy walks of brain-localized CD8 T cells observed previously [8]. One possibility is that Levy-like displacements arose due to special structures in the brain used by T cells for crawling. Such structures will then have to be poised at the threshold of percolation [57, 58]. Another possibility is that some brain-localized CD8 T cells are in fact located in brain vasculature and thus undergo Levy flights with the blood flow — similarly as we have observed for liver-localized CD8 T cells under some circumstances. Thirdly, T cells may be moving faster in places where other T cellshad left pathways, e.g., as has been proposed for T cells moving in collagen-fibrin gels [5]. Finally, Levy walks may arise in the analysis due to data generation or processing artifacts. We have found that when many T cells are present in the tissue it is sometimes difficult to segment experimental data to identify individual cells and their movements. We have also found in our and other publicly available datasets that cell coordinate data often have missing time frames, and if such tracks are not cleaned, data analysis may generate an impression of a long displacement between two assumed-to-be-sequential time frames. Whether such artifacts contributed to the overall detection of Levy walks of T cells in the brain remains to be determined, and in general it can be beneficial when raw data from imaging data analysis are made publicly available following publication (e.g., the data of Harris *et al.* [8] are not publicly available).

Our study has several limitations. Only a small area of the liver (approximately 500 × 500 × 50 *μ*m) is generally imaged with intravital microscopy which restricts the number of cells recorded and creates a bias for retention of slowly moving cells in longer movies. While we show that the digitized structure of liver sinusoids can override the intrinsic movement program of T cells, we did not estimate how much of the experimentally observed movement pattern of liver-localized T cells is due to a cell-intrinsic program such as a rare ability of cells to turn back, physical constraints of the liver sinusoids, or other environmental cues. (We did, however, found that even if Brownian-like walkers are not able to turn back in free 3D space, they superdiffuse for a relatively short time.) Methods for discriminating between these alternatives will have to be developed. Finally, we have focused on explaining our data with simplest possible movement models involving movement length distribution, MSD, and turning angles. Other, more complicated Levy walk models have been suggested such as generalized Levy walks (GLW) or Zigzag-GLW which allow for runs and pauses and assume or generate heterogeneity in cell velocities [8, 33, 38]. However, such models in general have more parameters, and fitting GLW-like models to our data is likely to result in overfitting. We could explain our data with relatively simple movement models such as correlated random walks or a combination of correlated random walks with Levy flights; Levy flights due to T cell floating in the blood were responsible for the fat-tailed distribution of movement lengths in our week 1 post RAS immunization data. Our results suggest that future studies focusing on benefits or evolutionary optimality of Levy walks would gain from more thorough investigation of the structural constraints and physiological tissue details that may be responsible for observed Levy-like movement pattern.

Our analyses raise many additional questions for future research. In particular, the impact of the *Plasmodium* infection on movement patterns of liver-localized CD8 T cells remains to be explored further. Our analysis of one unpublished dataset suggests similar long-term superdiffusion and short displacements for CD8 T cells following the infection (Fig. S5); however, this remains to be more fully investigated in the future. In particular, our recent work suggests that while initially, *Plasmodium*-specific CD8 T cells search for the infection site randomly, following formation of a T cell cluster near the parasite, other T cells become moderately attracted to the cluster [52, 54]. Whether movement patterns of liver-localized T cells change following formation of T cell cluster near the parasite remains to be investigated. Furthermore, infection may also influence blood flow allowing T cells in circulation to be recruited to the infection site. Therefore, it may be advantageous for T cells that are crawling on endothelial cells of the liver sinusoids to de-attach from the endothelium and rapidly float with the blood, thus, undergoing Levy flights. Whether a combination of local search by crawling and then large displacements with the blood flow allows for a more efficient localization of all parasites in the liver also remains to be determined.

## Abbreviations

MSD: mean squared displacement
LW: Levy walk
BW: Brownian walk
RAS: radiation attenuated sporozoites
FRC: fibroblastic reticular cells
DC: dendritic cells

## Ethics statement

All animal procedures were approved by the Animal Experimentation Ethics Committee of the Australian National University (Protocol numbers: A2016/17; 2019/36). All research involving animals was conducted in accordance with the National Health and Medical Research Council’s Australian Code for the Care and Use of Animals for Scientific Purposes and the Australian Capital Territory Animal Welfare Act 1992.

## Author contributions

VVG had the original idea for the study. VVG, HR, IAC, and JOC worked on various details of the experiments and analyses to be performed. JOC performed experiments (under supervision of IAC) and provided digitized data on movement of CD8 T cells in labeled liver sinusoids and separately provided images of labeled sinusoids. HR performed data curations, developed methods to digitize liver sinusoids and FRC networks and to analyze track data for T cells in the liver and LNs, ran most of data analyses and simulations, and wrote summaries of the methods and results. VVG wrote the first draft of the paper. All authors participated in the discussion of interim analyses and results. All authors contributed to the writing of the final version of the paper via changes, comments, and suggestions.

## Data sources

T cell movement/track data generated in this paper or in our previous publication are provided as an Excel spreadsheet with different tabs for different datasets. Imaging data of liver sinusoids (TIFF images) are also provided. The data from a previous publication on movement pattern of naive CD8 T cells in murine lymph nodes and imaging data of FRC networks in the LNs [46] are also provided. These files are stored together with the Matlab codes used to process and analyze the data on Github.

## Code sources

All analyses have been primarily performed in Matlab (ver 2019b) and codes used to generate most of figures in the paper are provided on GitHub: https://github.com/vganusov/Levy_walks_liver.git. All code questions should be addressed to H.R.

## Acknowledgments

We would like to thank Viktor Zenkov to contributing to the start of this work by manually digitizing liver sinusoidal maps and Marco Novkovic for providing imaging data of lymph nodes and data on movements of naive CD8 T cells in murine lymph nodes. This work was supported by the NIH grant (R01 GM118553) to VVG.

## Supplemental Information

### Additional movies, figures, and tables

**Movie S1:** Example of movement of liver-localized CD8 T cells with labeled liver sinusoids. Experiments were performed as described in Fig. 1 in 3 individual mice. In the movie red denotes rhodamine dextran-labeled liver sinusoids and green are *Plasmodium*-specific liver-localized CD8 T cells (OT-I). Cells turn yellow when there is an overlap between green and red. Bar scale is 50 *μ*m. The volume of 512 × 512 × 44 *μ*m was scanned every 13 sec.

**Figure S1:**
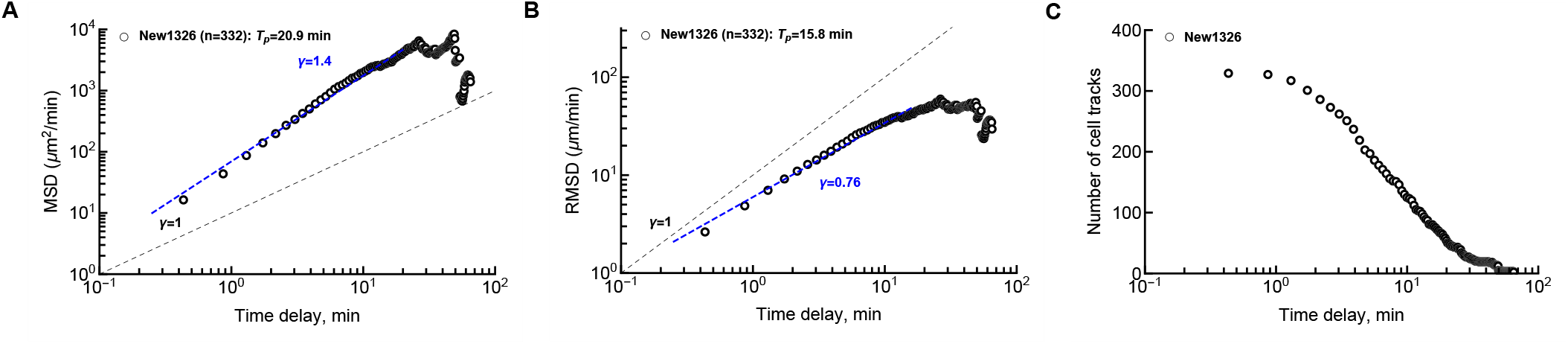
Departure from linear (on log-log scale) change in MSD or root mean squared displacement (RMSD) is in part due to decline in the number of cells with long tracks.√For the New1326 datasets of activated CD8 T cells in the liver we calculated the MSD (panel A), RMSD (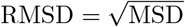, panel B), or the number of T cell tracks used to calculate these characteristics (panel C). Note that panel A lists a different slope from that given in Fig. 2A because different number of data points are taken into account in regression (*T*_*p*_ is the persistence time defining which data points were taken for linear regression).

**Table S1:** Estimates of parameters providing best fits of various datasets. We fitted Levy (eqn. (6)), Cauchy (eqn. (4)), General Pareto (eqn. (7)), exponential (eqn. (8)), or half-normal (eqn. (5)) to the data on movement of T cells in the liver, lymph nodes, or to branch length distribution data for liver sinusoids or fibroblastic retricular cell (FRC) network in lymph nodes. For distribution with a location parameter *r*_min_ we set *r*_min_ = 0. The same models were also fitted to scaled data (step length distribution data scaled by the RSMD). For imaging data we also list frequency of the imaging in the movies. See Extended Materials and Methods for the models and how models were fit to data.

| Non scaled data | Dataset | Levy $\gamma$ | Cauchy $\gamma$ | GP<br>k $\sigma$ | | Exp $\lambda$ | H-N $\sigma$ | Pareto<br>rmin ($\mu\text{m}$) μ-tail | | Sampling frequency (sec) |
| --- | --- | --- | --- | --- | --- | --- | --- | --- | --- | --- |
| Liver | New13 | 0.35 | 1.14 | -0.07 | 1.64 | 1.53 | 2.01 | 3.03 | 4.63 | 13.00 |
| Liver | New26 | 0.48 | 1.69 | 0.00 | 2.51 | 2.50 | 3.50 | 7.09 | 5.32 | 26.00 |
| Liver | New1326 | 0.49 | 1.69 | -0.01 | 2.51 | 2.49 | 3.46 | 7.09 | 5.47 | 26.00 |
| Liver | Wk1 | 0.13 | 0.40 | 0.28 | 0.50 | 0.76 | 2.15 | 0.61 | 2.55 | 6.50 |
| Liver | Wk4 | 0.10 | 0.33 | 0.01 | 0.46 | 0.46 | 0.66 | 0.94 | 4.42 | 3.25 |
| Liver | Lfa1_WT | 0.33 | 1.11 | 0.18 | 1.68 | 2.03 | 3.11 | 8.60 | 6.55 | 27.50 |
| Liver | Lfa1_KO | 0.18 | 0.57 | 0.11 | 0.80 | 0.91 | 1.46 | 2.10 | 3.30 | 27.50 |
| LN | 0209_Doc6 | 2.31 | 7.01 | -0.19 | 10.14 | 8.72 | 10.58 | 22.02 | 6.67 | 20.00 |
| LN | 0409_Doc7 | 1.20 | 3.67 | -0.48 | 5.77 | 4.16 | 4.71 | 6.14 | 6.74 | 20.00 |
| Scaled data | Dataset | Levy $\gamma$ | Cauchy $\gamma$ | General Pareto<br>k $\sigma$ | | Exp $\lambda$ | H-N $\sigma$ | Pareto<br>ρmin μ-tail | | Sampling frequency (sec) |
| Liver | New13 | 0.23 | 0.74 | -0.07 | 1.07 | 1.00 | 1.31 | 2.01 | 4.68 | 13.00 |
| Liver | New26 | 0.19 | 0.68 | 0.00 | 1.01 | 1.00 | 1.41 | 2.90 | 5.39 | 26.00 |
| Liver | New1326 | 0.20 | 0.68 | -0.01 | 1.01 | 1.00 | 1.39 | 2.91 | 5.55 | 26.00 |
| Liver | Wk1 | 0.17 | 0.52 | 0.28 | 0.66 | 1.00 | 2.82 | 0.78 | 2.52 | 6.50 |
| Liver | Wk4 | 0.22 | 0.71 | 0.01 | 0.99 | 1.00 | 1.44 | 1.93 | 4.28 | 3.25 |
| Liver | Lfa1_WT | 0.16 | 0.55 | 0.18 | 0.83 | 1.00 | 1.53 | 4.42 | 6.47 | 27.50 |
| Liver | Lfa1_KO | 0.20 | 0.63 | 0.11 | 0.88 | 1.00 | 1.60 | 2.29 | 3.30 | 27.50 |
| LN | 0209_Doc6 | 0.29 | 0.87 | -0.38 | 1.30 | 1.00 | 1.14 | 2.05 | 9.34 | 20.00 |
| LN | 0409_Doc7 | 0.29 | 0.88 | -0.48 | 1.39 | 1.00 | 1.13 | 1.55 | 7.00 | 20.00 |
| Structure | Dataset | Levy $\gamma$ | Cauchy $\gamma$ | GP<br>k $\sigma$ | | Exp $\lambda$ | H-N $\sigma$ | Pareto<br>rmin ($\mu\text{m}$) μ-tail | | |
| Liver sinusoids | Evans Blue | 3.48 | 10.54 | -0.20 | 15.71 | 13.38 | 16.37 | 36.36 | 7.45 |  |
| LN FRCs | CCL19day5 | 2.31 | 7.01 | -0.19 | 10.14 | 8.72 | 10.58 | 22.02 | 6.67 |  |

**Table S2:** AIC values and Akaike weights of alternative models fit to different datasets (as described in Tab. S1). See Extended Materials and Methods for the models and how models were fit to data.

| Non scaled data | Dataset | Levy | Cauchy | GP | Exp | H-N | Levy | Cauchy | GP | Exp | H-N |
| --- | --- | --- | --- | --- | --- | --- | --- | --- | --- | --- | --- |
| Liver | New13 | 11397 | 11433 | 7269 | 7302 | 7279 | 0.00 | 0.00 | 0.99 | 0.00 | 0.01 |
| Liver | New26 | 40929 | 41951 | 29623 | 29621 | 30631 | 0.00 | 0.00 | 0.32 | 0.68 | 0.00 |
| Liver | New1326 | 45728 | 46801 | 33002 | 33003 | 33968 | 0.00 | 0.00 | 0.58 | 0.42 | 0.00 |
| Liver | Wk1 | 5868 | 5968 | 2746 | 3398 | 6955 | 0.00 | 0.00 | 1.00 | 0.00 | 0.00 |
| Liver | Wk4 | 8184 | 8275 | 1860 | 1859 | 2575 | 0.00 | 0.00 | 0.39 | 0.61 | 0.00 |
| Liver | Lfa1_WT | 26909 | 28181 | 19649 | 19789 | 21556 | 0.00 | 0.00 | 1.00 | 0.00 | 0.00 |
| Liver | Lfa1_KO | 15307 | 15615 | 8554 | 8668 | 10549 | 0.00 | 0.00 | 1.00 | 0.00 | 0.00 |
| LN | 0209_Doc6 | 42313 | 41380 | 28946 | 30515 | 28685 | 0.00 | 0.00 | 0.00 | 0.00 | 1.00 |
| LN | 0409_Doc7 | 3352 | 3278 | 2281 | 2433 | 2281 | 0.00 | 0.00 | 0.53 | 0.00 | 0.47 |
| Scaled data | Dataset | AIC |  |  |  |  | Akaike Weights |  |  |  |  |
|  |  | Levy | Cauchy | GP | Exp | H-N | Levy | Cauchy | GP | Exp | H-N |
| Liver | New13 | 9219 | 9254 | 5090 | 5123 | 5099 | 0.00 | 0.00 | 0.99 | 0.00 | 0.01 |
| Liver | New26 | 26806 | 27820 | 15491 | 15489 | 16494 | 0.00 | 0.00 | 0.32 | 0.68 | 0.00 |
| Liver | New1326 | 30021 | 31086 | 17287 | 17287 | 18248 | 0.00 | 0.00 | 0.59 | 0.41 | 0.00 |
| Liver | Wk1 | 7141 | 7242 | 4020 | 4673 | 8231 | 0.00 | 0.00 | 1.00 | 0.00 | 0.00 |
| Liver | Wk4 | 14498 | 14595 | 8181 | 8180 | 8901 | 0.00 | 0.00 | 0.41 | 0.59 | 0.00 |
| Liver | Lfa1_WT | 18723 | 19988 | 11456 | 11596 | 13359 | 0.00 | 0.00 | 1.00 | 0.00 | 0.00 |
| Liver | Lfa1_KO | 16191 | 16499 | 9439 | 9554 | 11435 | 0.00 | 0.00 | 1.00 | 0.00 | 0.00 |
| LN | 0209_Doc6 | 24765 | 23829 | 11394 | 12965 | 11133 | 0.00 | 0.00 | 0.00 | 0.00 | 1.00 |
| LN | 0409_Doc7 | 1924 | 1849 | 853 | 1005 | 853 | 0.00 | 0.00 | 0.54 | 0.00 | 0.46 |
| Structure | Dataset | Levy | Cauchy | GP | Exp | H-N | Levy | Cauchy | GP | Exp | H-N |
| Liver sinusoids | Evans Blue | 47544 | 47114 | 37936 | 38375 | 37599 | 0.00 | 0.00 | 0.00 | 0.00 | 1.00 |
| LN FRCs | CCL19day5 | 29759 | 29402 | 23058 | 23349 | 22758 | 0.00 | 0.00 | 0.00 | 0.00 | 1.00 |

**Figure S2:**
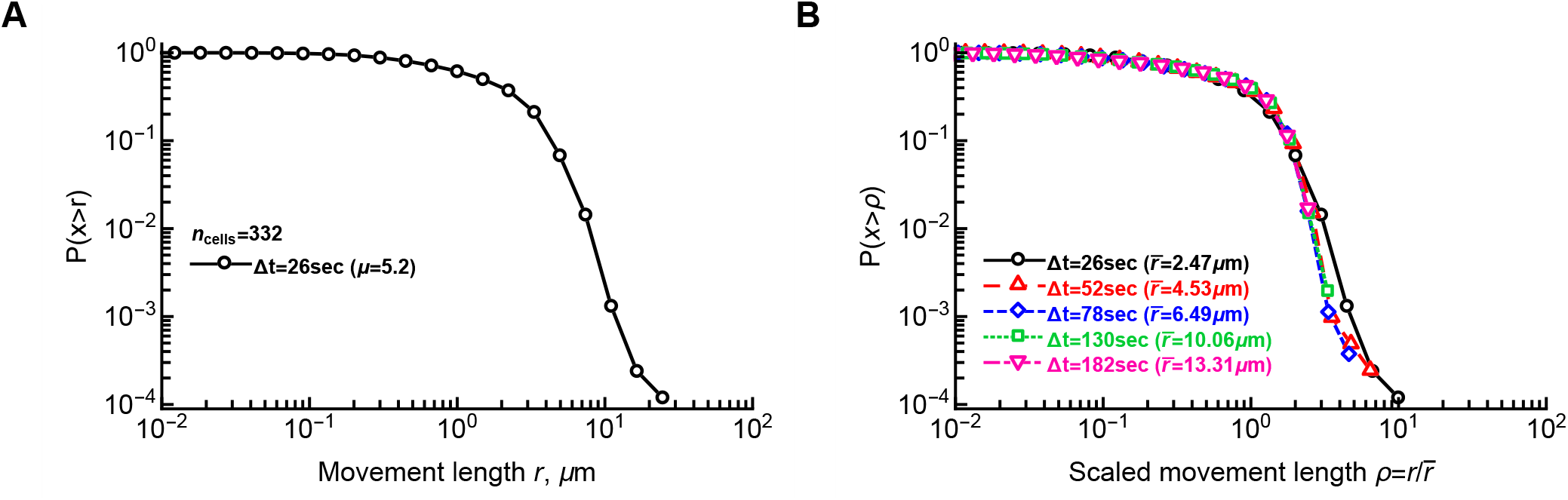
Liver-localized display similar displacement distribution for sub-sampled data. For the cleaned New1326 dataset we calculated the cumulative distribution of displacements for all data (panel A) or when we sub-sample the data, i.e., by taking every time frame (Δ*t* = 26 sec), every second time frame (Δ*t* = 52 sec), etc (panel B). In panel B we rescaled all displacements by the average displacement 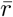 in a given sub-sampled dataset. Results were similar if we scaled the data by dividing displacements by the relative duration of the sub-sampling period (*k* = 1, 2 ...). Parameter *μ* in panel A is from the fit of the Pareto distribution (with *r*_min_ = 9.1 *μ*m) to the subset of the data (fail fit in which data with *r* ≥ *r*_min_ were used).

**Figure S3:**
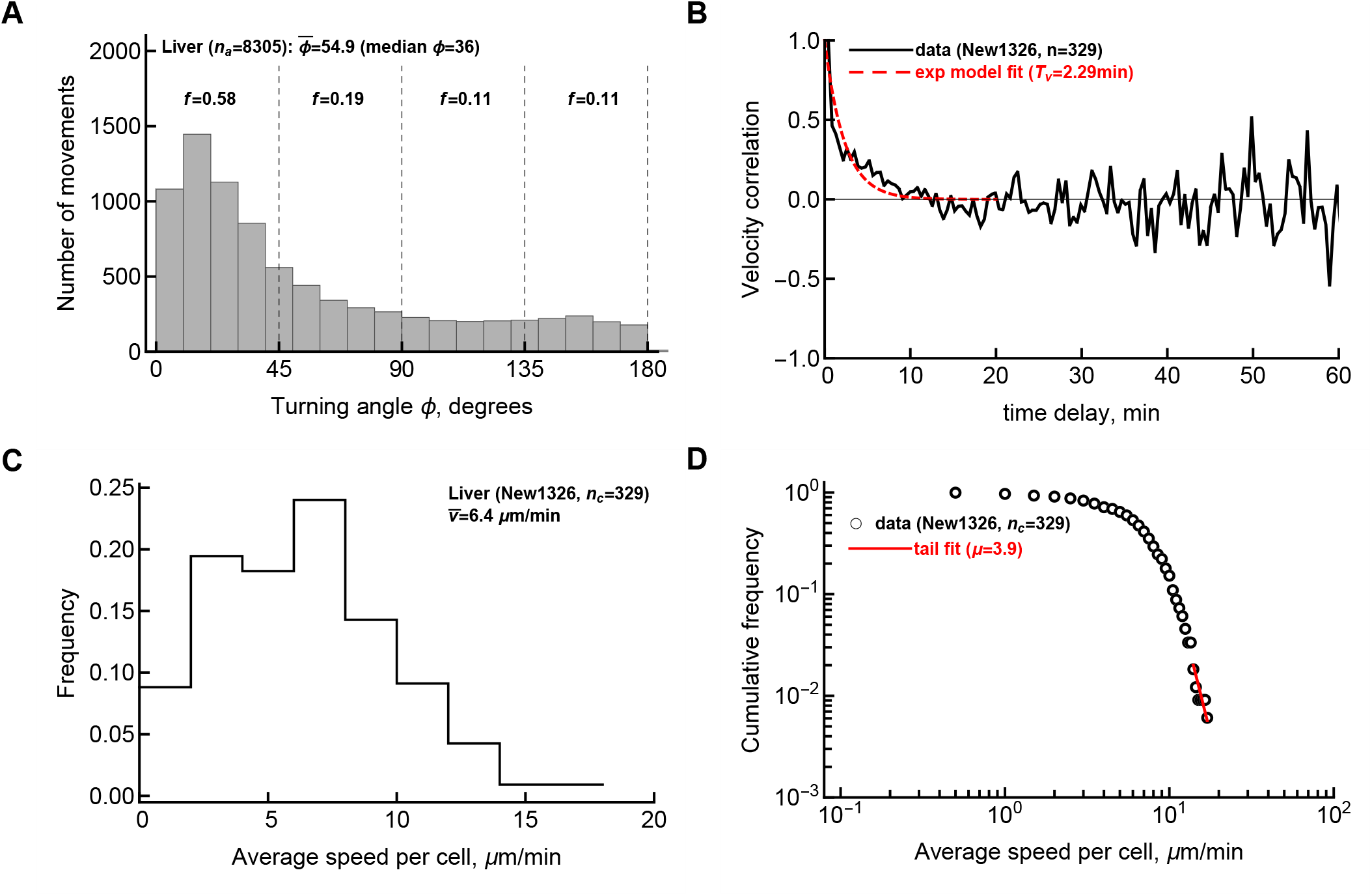
Activated CD8 T cells in the liver perform correlated random walk with a high degree of persistence. A: For every track in our dataset that includes measurement of T cell movement in the liver at 26 sec time steps, we calculated the turning angles for each subsequent cell movement. The data are from 3 individual mice involving 332 cleaned tracks and 8305 movements (“New1326” dataset). Fraction *f* denotes the frequency of turning angles between specific cutoff values. B: We calculated the correlation between vectors determining the first movement and every another movement for all 329 cell tracks in our data for which velocity correlations could be calculated. Specifically, we calculated the angle *ϕ* between the vector determining the first cell movement and every another movement at different times and plotted cos(*ϕ*) vs. the time difference between first movement and another movements. Insert in B shows the data for the first 20 min of imaging. The exponential decay model (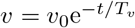 was fitted to the data and estimated parameter *T*_*v*_ is shown. C: Distribution of average speed per cell in these data. D: cumulative distribution of speeds of all cells int he population. Parameter *μ* denotes the shape parameter of the Pareto distribution fitted to the tail of these data.

**Figure S4:**
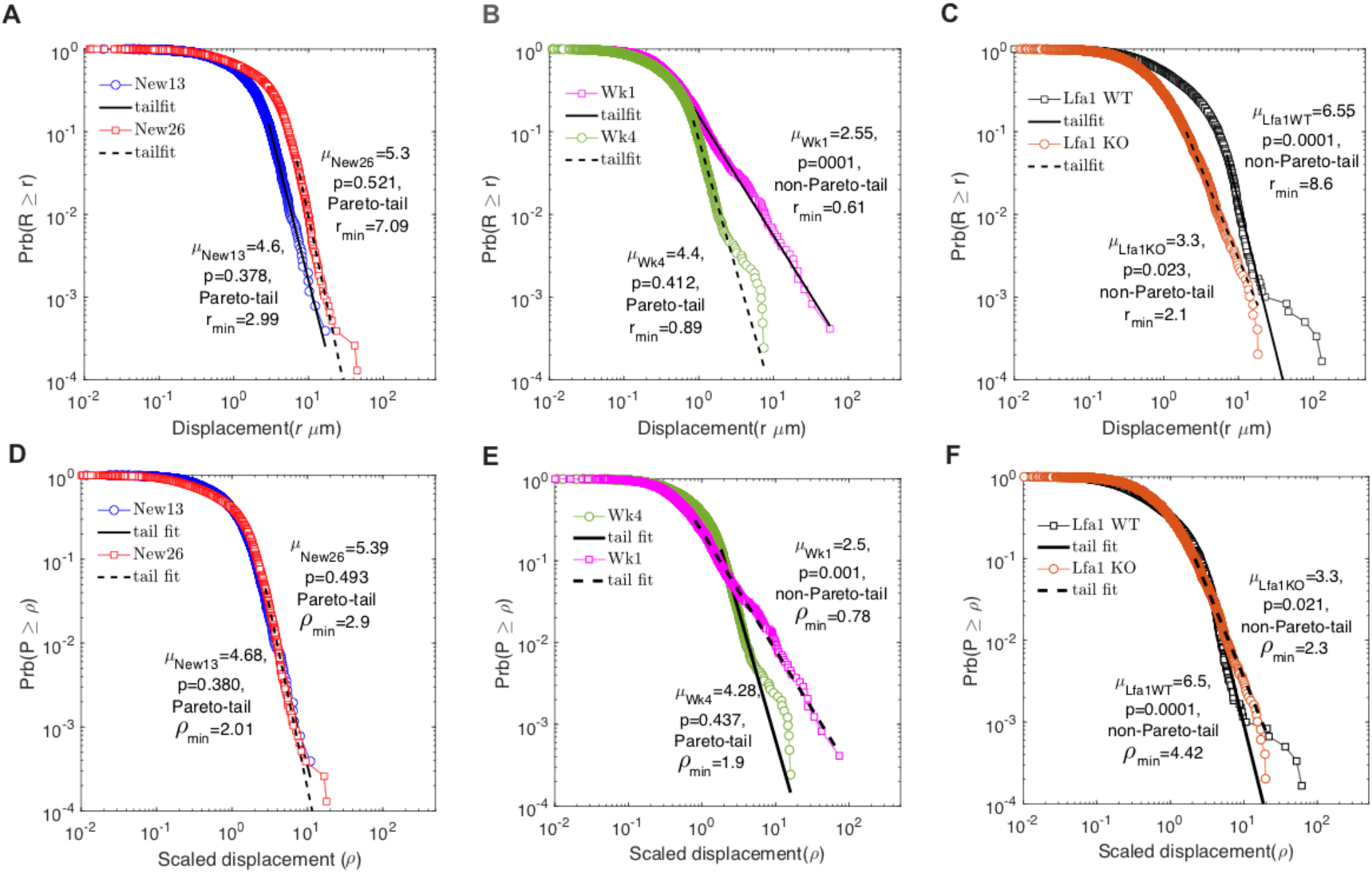
Fits of the Pareto (powerlaw) distributions to the CD8 T cell in the liver displacement data. We used previously published method [34] to estimate the shape parameter *μ* of the Pareto distribution (eqn. (9)) for 6 different datasets (see Materials and Methods for dataset description) by fitting a linear function (on log-log scale) to the tails of the data distribution using likelihood method. We fit the model to actual displacements *r* (A-C) or to scaled displacements *ρ* (D-F). The cut-off *r*_min_ (for actual displacements) or *ρ*_min_ (for normalized displacements) for the linear regressions was chosen using a goodness-of-fit based method [34]. In short, for multiple choices of *r*_min_ slope *μ* was estimated via the method of maximum likelihood and the Kolmogorov-Smirnov goodness of fit statistic, *D*, was calculated. Value of *r*_min_ that resulted in minimal fit statistics *D* was chosen for the final fit. Indicated *p* values were estimated by resampling the data 1000 times and performing Kolmogorov-Smirnov Goodness of Fit Test [34].

**Figure S5:**
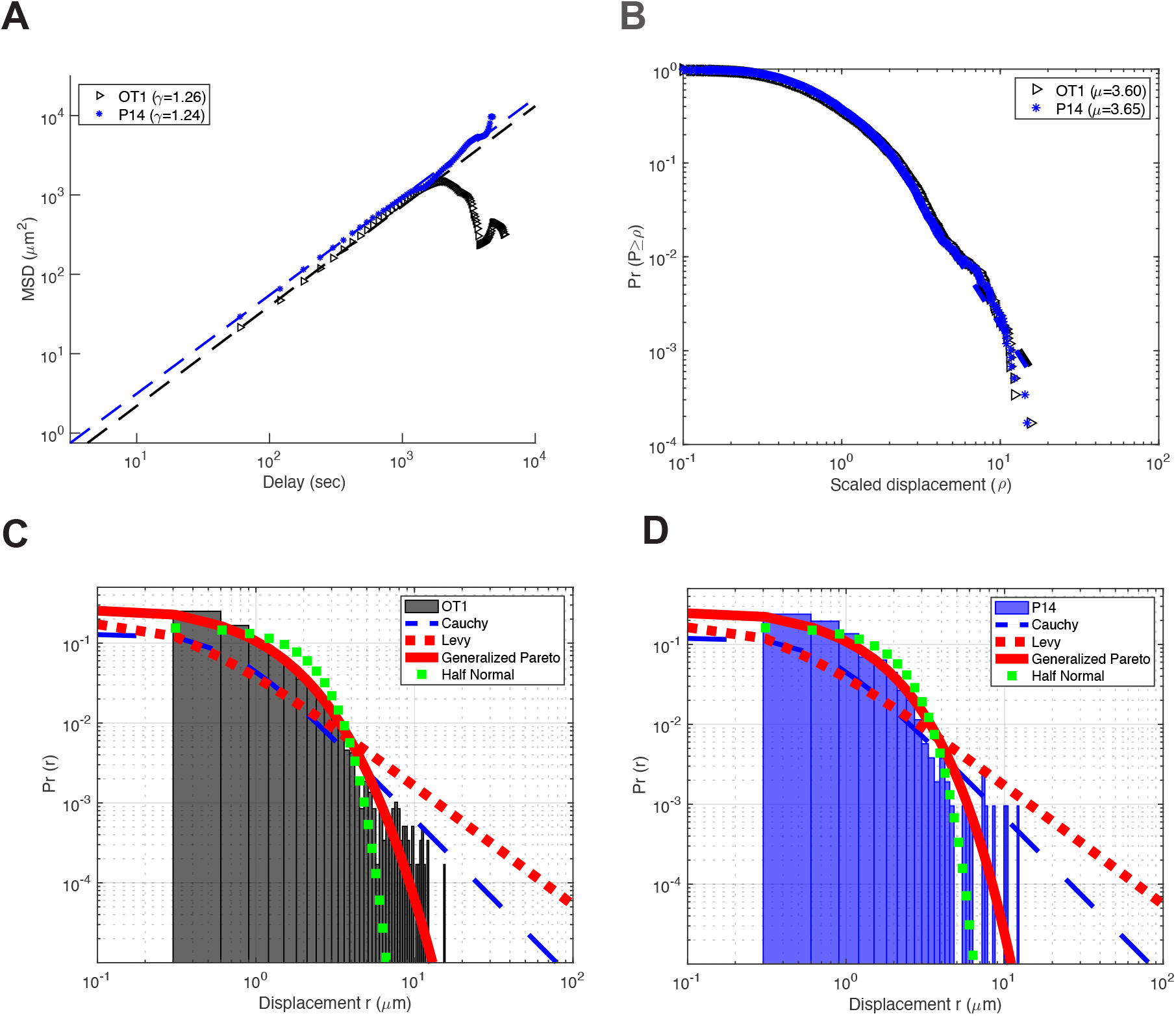
Activated CD8 T cells are superdiffusive Brownian walkers in the liver during *Plasmodium* infection. We performed experiments in which *in vitro* activated OT-I (*Plasmodium berghei*-specific) and P14 (LCMV-specific) CD8 T cells were transferred i.v. into naive mice and these mice were 1.5-2 hours later infected with 10^5^ *Plasmodium berghei*-CS^5M^ sporozoites expressing SIINFEKL epitope from chicken ovalbumin [31, 62]. Ten to twenty minutes after infection, livers of infected mice were exposed and movements of T cells were recorded using intravital microscopy [52]. We performed the same analyses as in the main text (Fig. 2) including (A) regression analysis of the MSD with time, (B) distribution of movement lengths, (C)-(D) fitting alternative distributions to the movement data for OT1 (C) or P14 (D) T cells. Generalized Pareto distribution was best describing data in C-D (results not shown).

**Figure S6:**
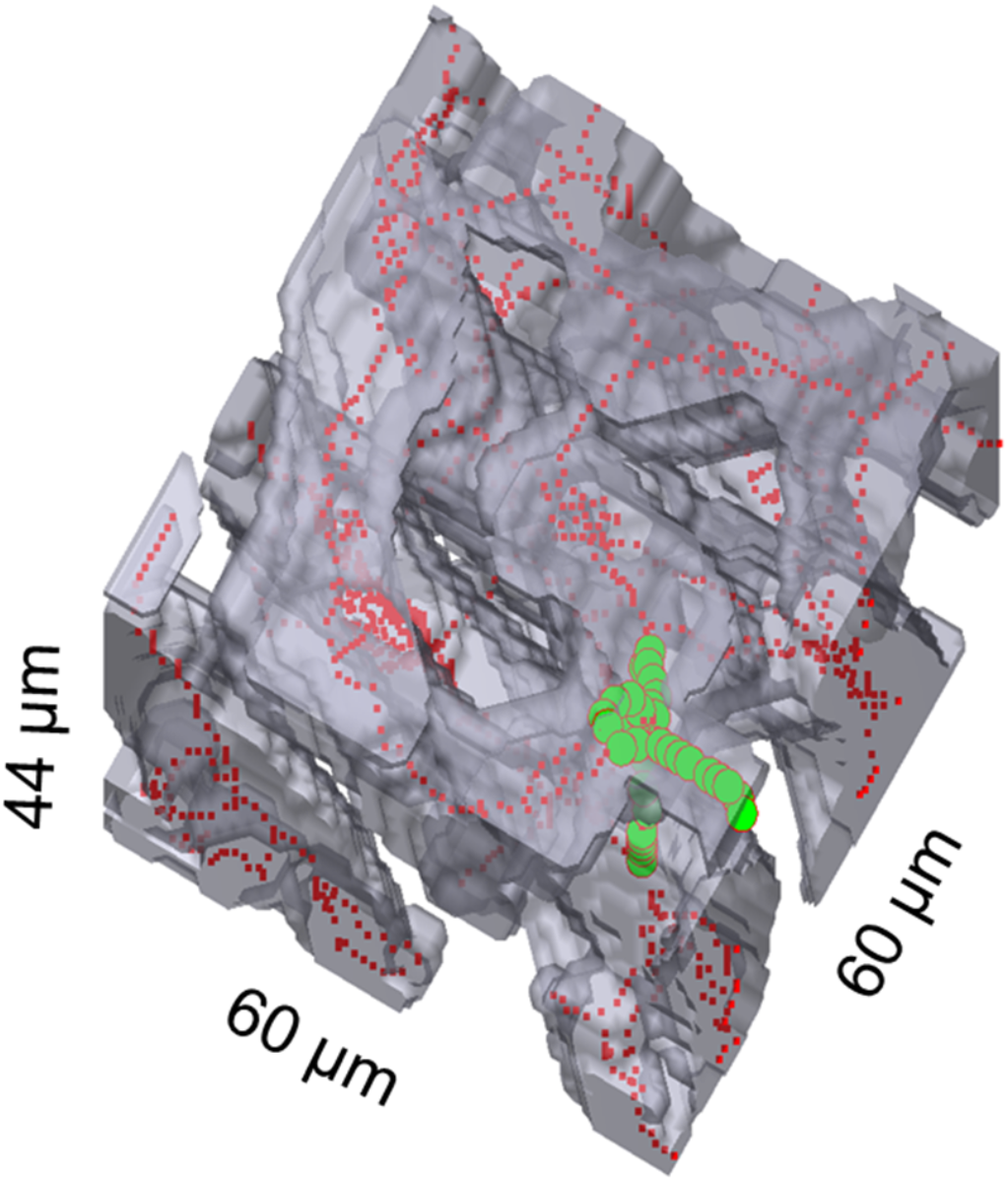
Example of the simulation in which T cells were moving in sinusoids of the liver. Grey denotes sinusoids, red dots denote the mean path of the sinusoids, and green denotes moving T cell.

**Figure S7:**
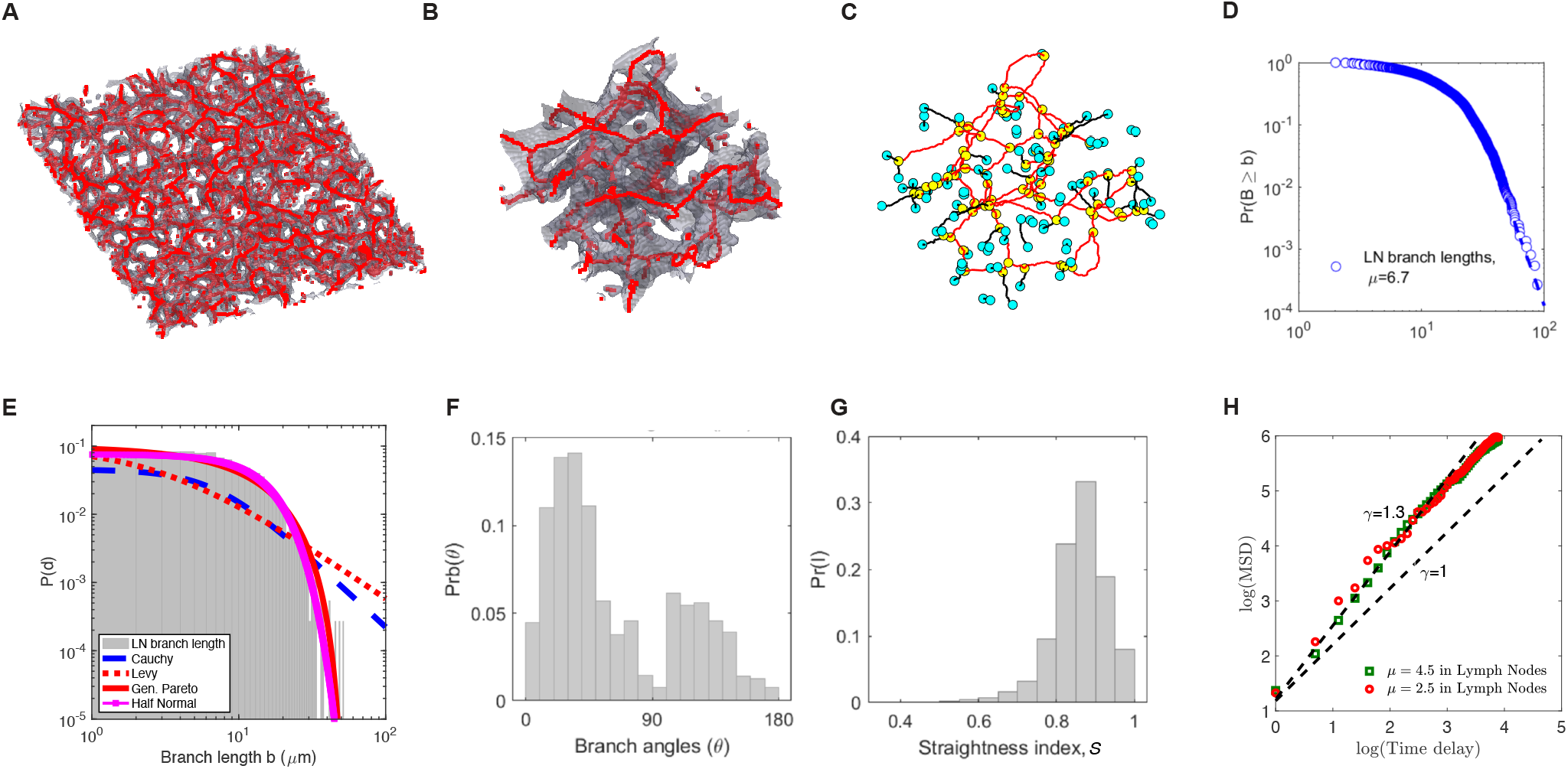
Characterizing structure of the fibroblastic reticular cell network in the murine lymph nodes. We used previously published data in which FRCs were imaged using Ccl19^eyfp^ reporter mouse [46]. As with analysis of liver sinusoids z-stacks of LN images were contrasted and mean paths in the FRCs were calculated. To simulate T cell walk on digitized FRC network we assumed that movement length distribution is described by the powerlaw (Pareto) distribution with *r*_min_ = 0.2 *μ*m and *μ* = 4.5 (for Brownian walks) or *μ* = 2.5 (for Levy walks).

**Figure S8:**
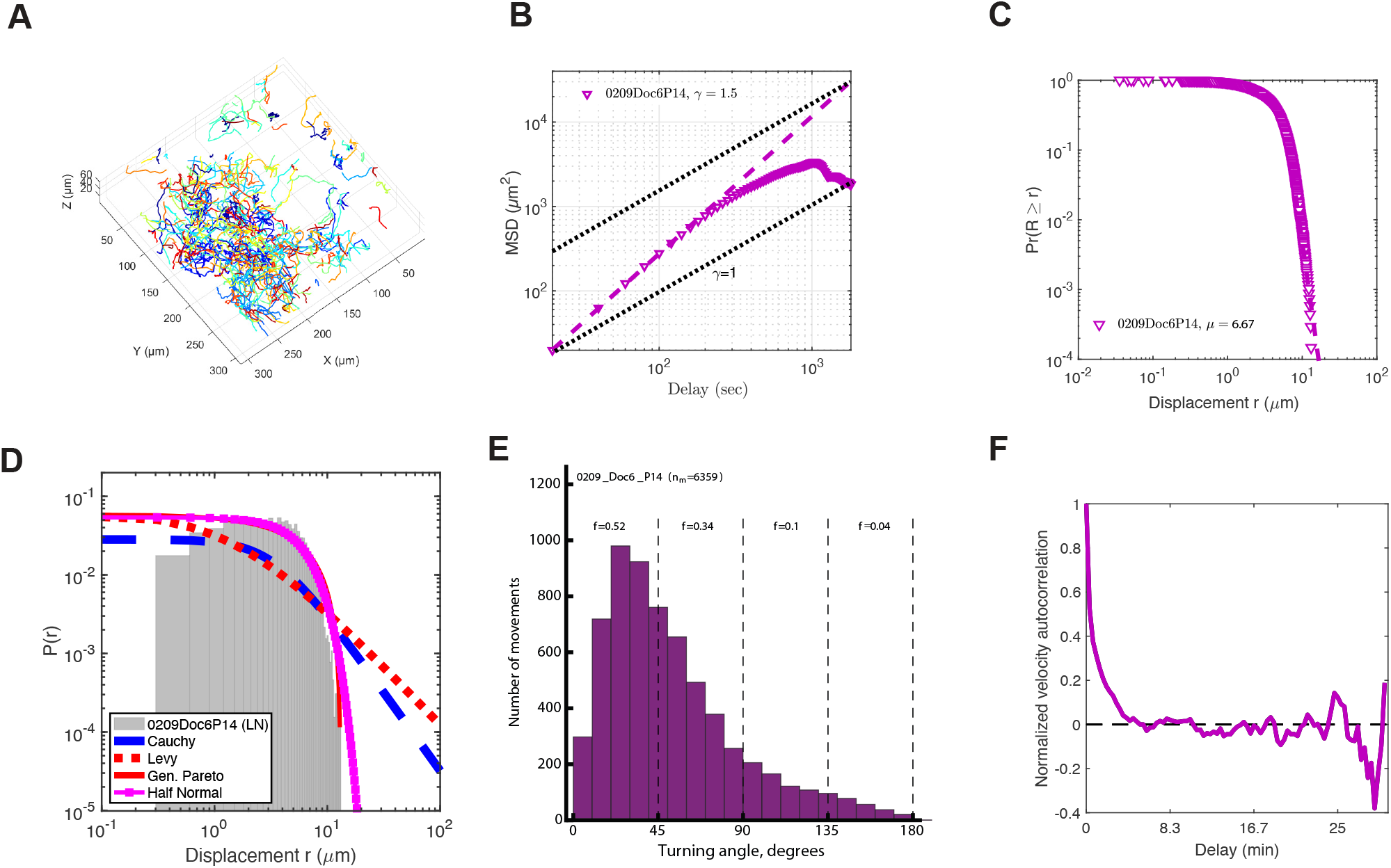
Naive CD8 T cells in lymph nodes are not superdiffusive. We analyzed experimental data in which naive CD8 T cells moving in resting lymph nodes were imaged using two photon microscope [46]. Example of the analysis of data from one of 6 such datasets is show. A: individuals tracks of T cells digitized using Imaris software; B: MSD vs. time for all T cels indicating short-term (2-3 min) persistence/superdiffusion; C: movement length distribution with tail analysis performed [34]; D: fits of several alternative distributions to the movement length distribution data; E: distribution of turning angles of T cells in the LNs; F: velocity autocorrelation function for T cells moving in the LNs.

**Figure S9:**
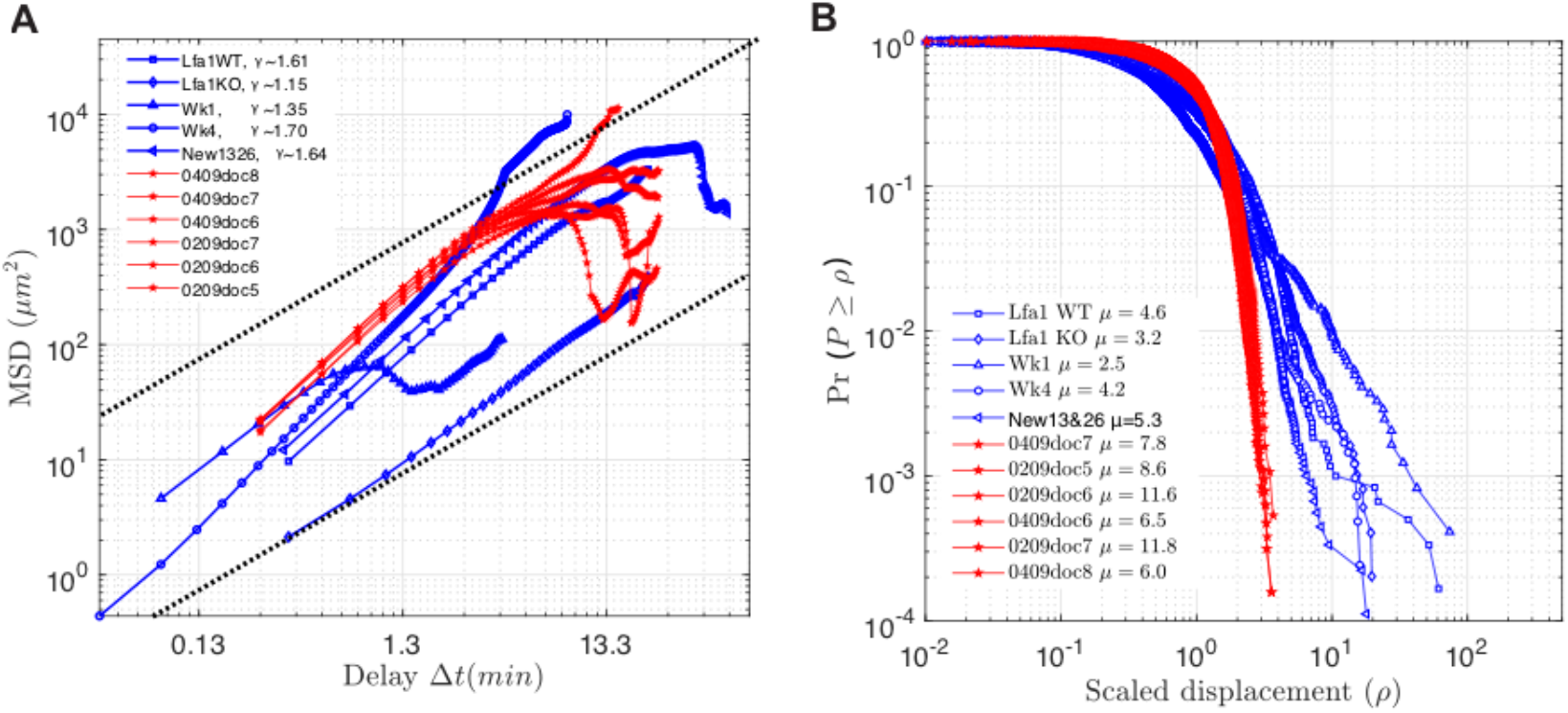
T cells in the liver display persistent movement for a longer time as compared to T cells in the LNs. Data from all datasets on movement of activated CD8 T cells in the liver (blue) or naive CD8 T cells in the LNs (red) are plotted together. A: difference in MSD for T cells in the LNs vs. liver; B: difference in movement length distribution of T cells in the liver vs. LNs.

